# Identification of a gene expression signature of vascular invasion and recurrence in stage I lung adenocarcinoma via bulk and spatial transcriptomics

**DOI:** 10.1101/2024.06.07.597993

**Authors:** Dylan Steiner, Lila Sultan, Travis Sullivan, Hanqiao Liu, Sherry Zhang, Ashley LeClerc, Yuriy O. Alekseyev, Gang Liu, Sarah A. Mazzilli, Jiarui Zhang, Kimberly Rieger-Christ, Eric J. Burks, Jennifer Beane, Marc E. Lenburg

## Abstract

Microscopic vascular invasion (VI) is predictive of recurrence and benefit from lobectomy in stage I lung adenocarcinoma (LUAD) but is difficult to assess in resection specimens and cannot be accurately predicted prior to surgery. Thus, new biomarkers are needed to identify this aggressive subset of stage I LUAD tumors. To assess molecular and microenvironment features associated with angioinvasive LUAD we profiled 162 resected stage I tumors with and without VI by RNA-seq and explored spatial patterns of gene expression in a subset of 15 samples by high-resolution spatial transcriptomics (stRNA-seq). Despite the small size of invaded blood vessels, we identified a gene expression signature of VI from the bulk RNA-seq discovery cohort (n=103) and found that it was associated with VI foci, desmoplastic stroma, and high-grade patterns in our stRNA-seq data. We observed a stronger association with high-grade patterns from VI^+^ compared with VI^-^ tumors. Using the discovery cohort, we developed a transcriptomic predictor of VI, that in an independent validation cohort (n=60) was associated with VI (AUROC=0.86; p=5.42×10^-6^) and predictive of recurrence-free survival (HR=1.98; p=0.024), even in VI^-^ LUAD (HR=2.76; p=0.003). To determine our VI predictor’s robustness to intra-tumor heterogeneity we used RNA-seq data from multi-region sampling of stage I LUAD cases in TRACERx, where the predictor scores showed high correlation (R=0.87, p<2.2×10^-16^) between two randomly sampled regions of the same tumor. Our study suggests that VI-associated gene expression changes are detectable beyond the site of intravasation and can be used to predict the presence of VI. This may enable the prediction of angioinvasive LUAD from biopsy specimens, allowing for more tailored medical and surgical management of stage I LUAD.

## INTRODUCTION

Lung adenocarcinoma (LUAD) is the most common lung cancer subtype and invasive LUAD represents 70-90% of surgically resected lung cancers^1^. Improved early detection through computed tomography (CT)-based screening programs is expected to increase the proportion of LUAD diagnosed at an early stage. Microscopic vascular invasion (VI), defined as tumor invasion within the lumen of veins or arteries, is a well described route to metastatic dissemination and is consistently associated with higher rates of tumor recurrence among stage I LUAD^2–7^. VI is not included in the current World Health Organization (WHO) grading system for lung cancer, but may be a better predictor of recurrence than the most severe WHO-2021 grade, leading to our proposal for VI^+^ LUAD to be reclassified as distinct angioinvasive LUAD^8^.

Patients with stage I VI+ LUAD may benefit from adjuvant therapy^9–12^, but it is difficult to assess in resected tumor specimens. Elastic stains can be used to improve visualization of invaded blood vessels over hematoxylin and eosin (H&E), but comprehensive tumor histopathology review for small (<1mm) VI foci is difficult and prone to false negatives. Similarly, biopsy specimens do not provide enough tissue material to identify VI^+^ LUAD prior to surgery, preventing the use of VI status to guide surgical or neoadjuvant treatment approaches.

Recent evidence from large clinical trials including JCOG0802/WJOG4607L and CALGB140503 suggests that lobectomy provides no clinical benefit over sublobar resection for patients with stage IA disease^13,14^, although post-hoc analyses are beginning to show subsets of patients for which lobectomy might decrease the rate of recurrence^15^. Moreover, retrospective analysis shows that VI^+^ patients receiving sublobar resection have increased rates of recurrence, highlighting an emerging need to identify if these patients will benefit from precision surgical approaches^16^.

Molecular profiling technologies are common in clinical pathology but prior RNA-sequencing (RNA-seq) studies of LUAD have primarily focused on identifying signatures of outcome, which lack reproducibility^17,18^. Others have identified signatures associated with high-grade growth patterns or subtypes^19,20^, such as solid LUAD. To our knowledge, VI-associated gene expression has not been studied. Pathologists infrequently document VI as a separate entity from lymphatic invasion (LI), preferring to designate the presence of either type as lymphovascular invasion (LVI). This obfuscates the independent prognostic value of VI^21^ and complicates downstream molecular approaches seeking to disentangle these modes of tumor spread, the differences of which are still incompletely understood^22^. New advances in spatial transcriptomics (stRNA-seq) allow for resolving transcriptomic changes associated with tumor invasion within the spatial context of the tumor and tumor microenvironment^23^. This represents an opportunity to elucidate VI^+^ LUAD biology with the aim to improve detection of these tumors. In this study, we profiled 163 tumors with detailed histopathologic annotations by RNA-seq, including 15 by stRNA-seq, from a diverse multi-institutional cohort of stage I LUAD patients to comprehensively define the transcriptional landscape of angioinvasive LUAD. We derived a novel molecular signature associated with VI^+^ LUAD using bulk RNA-seq, described its association with histopathology *in situ* using stRNA-seq, developed and validated a machine learning based predictor of VI and demonstrated its robustness to LUAD intra-tumor heterogeneity (ITH).

## RESULTS

### VI is the invasion type most associated with stage I LUAD recurrence

We assembled a retrospective discovery cohort of stage I LUAD tumors (n=192) from patients across two institutions, Boston Medical Center (BMC), a large urban safety-net hospital, and Lahey Hospital and Medical Center (LHMC), a suburban hospital, a subset of a previously published cohort with detailed clinicopathology annotations^8^. Tumors were graded using our novel grading system, which classifies tumors into low malignant potential (LMP), no special type (NST), or VI, is associated more associated with outcome than the World Health Organization (WHO) grading system (Extended Data Fig. 1a-b) and whose reproducibility has been validated by us and others^8,16,24,25^. VI was the strongest predictor (hazard ratio (HR)=7.62, 95% CI 3.70-15.8, Cox regression p=5.17×10^-8^) of 7-year recurrence-free survival (RFS) among the four observed modes of tumor spread (VI, LI, visceral pleural invasion – VPI, and spread through air spaces – STAS), in agreement with previous reports of its prognostic value in stage I LUAD^8,26–28^ (Extended Data Fig. 1c-f). VI was present in 30% of tumors, and 84% of VI^+^ tumors contained multiple types of invasion (Extended Data Fig. 1g). Regardless, VI remained a significant predictor of recurrence in when including clinical covariates and the other invasion types (HR = 11.1, 95% CI 3.86–32.3, multivariate Cox regression p=8.59×10^-6^; Extended Data Fig. 1h).

When scoring tumors based on the proportion of LUAD growth patterns, VI was associated with solid (Bonferroni adjusted P value (p.adj) = 0.004, Wilcoxon test), but not micropapillary (p.adj=0.08) or cribriform proportion (p.adj=0.35) (Extended Data Fig. 1i). In contrast, STAS was most strongly positively associated with increased micropapillary proportion (p.adj=5.3 x10^-7^), as previously reported.^29^ VI^+^ tumors were more strongly associated with distant recurrence (sub-distribution HR=19.1, 95% CI 4.27–85.0, multivariable Fine-Gray regression p=1.1×10^-4^) than loco-regional recurrence only (sub-distribution HR=3.86, 95% CI 1.22–12.2, multivariable Fine-Gray regression p= 0.02), consistent with tumor spread through the vascular system (Extended Data Figs. 1j and k).

### Gene expression changes in stage I LUAD with VI

To identify gene expression changes in tumors with VI, we profiled a subset of surgically resected stage I LUAD tumors from the discovery cohort (n = 108, n = 103, post-QC) by bulk RNA-sequencing (RNA-seq) (Fig. 1a, Table S1). Among tumors that passed quality control, 78 were VI^-^ and 25 were VI^+^. We identified 474 genes differentially expressed between VI^+^ tumors and tumors of low malignant potential (LMP) using the three-level factor of tumor grade (LMP, NST, VI) as the independent variable at a significance threshold of FDR < 0.01, which we clustered into four gene expression clusters via hierarchical clustering (Fig. 1b). Genes in three of the clusters had increased expression in LUAD with VI (clusters 1, 2 and 3) and genes in the other cluster had decreased expression (cluster 4). Pathway enrichment analysis using EnrichR (FDR < 0.05) revealed that genes involved in different biological processes were enriched in each cluster (Fig. 1c). Cluster 1 (n=115 genes) was enriched for genes involved in cell cycle. In contrast, cluster 2 (n=37 genes) was enriched for genes involved in tissue remodeling and vasculogenesis, including the epithelial to mesenchymal transition (EMT), extracellular matrix organization, and angiogenesis. The genes in cluster 3 (n=182 genes) were enriched for some pathways that overlapped with cluster 1 like mTORC1 signaling and E2F targets but was distinguished by enrichment for pathways related to response to reactive oxygen species and hypoxia. Finally, the genes with decreased expression in VI tumors (cluster 4; n=140 genes) were enriched for pathways indicating reduced malignant progression (p53 pathway, regulation of cell growth), increased cell-cell adhesion via TGF-β signaling (regulation of pathway-restricted SMAD protein phosphorylation) and increased immune surveillance (IL-2/STAT5 signaling) in VI tumors. In the discovery cohort, the mean z-score of the genes in each of the four clusters was a significant predictor of VI (Extended Data Fig. 2a) and was also able to distinguish between NST and VI^+^ tumors (Extended Data Fig. 2b), supporting that the differentially expressed genes reflect VI-specific biology. Given that VI and LI are routinely reported together as LVI^8,21^, we compared the VI signature with gene expression differences associated with LI in stage I LUAD. When we added LI as a covariate to our model, we identified 133 genes associated with VI (FDR < 0.01), which included 130 genes from the original 474-gene VI signature (Extended Data Fig. 3a). At the same significance threshold (FDR < 0.01), we recovered only 6 genes associated with LI, with 2/6 belonging to the VI signature (Extended Data Fig. 3b), suggesting the biological signal for LI is much weaker. To further interrogate the relationship between LI and VI associated gene expression we explored whether the various VI gene clusters from the 474-gene signature are enriched among the genes most differentially expressed in LI+ tumors using gene-set enrichment analysis (GSEA). Interestingly, we found that the genes in VI Cluster 4 are significantly enriched among the genes decreased in LI+ tumors. And unlike the genes in VI Cluster 1 (cell-cycle) and VI Cluster 3 (hypoxia), the genes in VI Cluster 2 (EMT/angiogenesis), were not significantly enriched among the genes most increased in LI^+^ tumors (Extended Data Fig. 3c-f).

**Figure 1.**
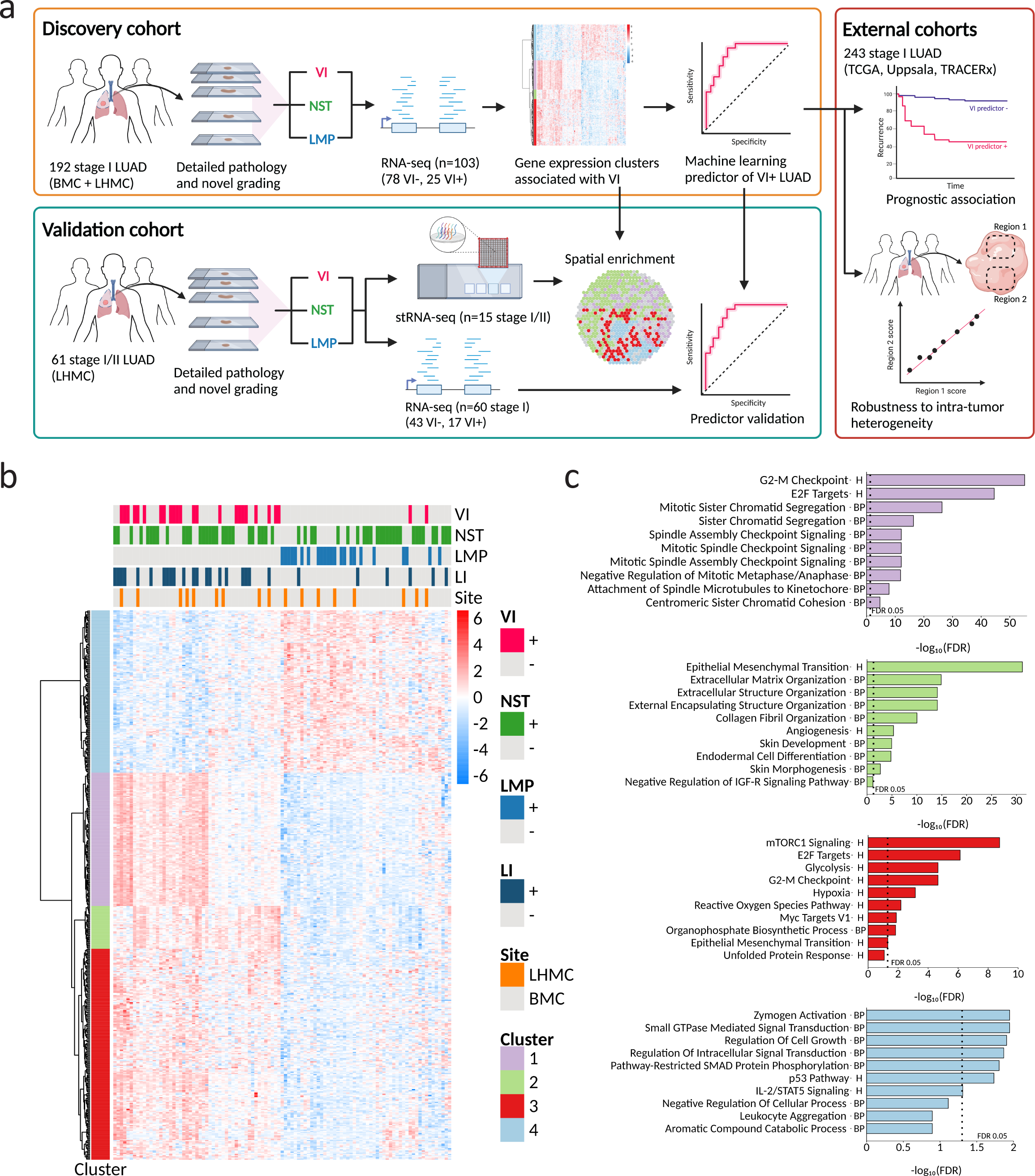
Study overview and distinct gene expression changes associated with VI in stage I LUAD. **a**. Overview of cohorts, sequencing technologies, and analyses utilized in the study. VI, vascular invasion; NST, no special type; LMP, low malignant potential; BMC, Boston Medical Center; LHMC, Lahey Hospital and Medical Center. All tumors were selected to be stage I at the time of collection except for one tumor profiled by stRNA-seq that was upstaged to stage IIA under the 8^th^ TNM edition. **b**. Co-expression heatmap of 474 genes differentially expressed between VI and LMP (FDR < 0.01) in a discovery cohort of stage I LUAD tumors (n=103) grouped into k=4 clusters. Heatmap units are log counts per million (CPM) scaled by transcript. LI, lymphatic invasion. **c**. Top 10 Biological enrichment terms of genes within each gene co-expression cluster. Pathways were ranked within each cluster by FDR values obtained from inputting cluster genes into the EnrichR database with queries for MSigDB Hallmark 2020 (H) and GO Biological Process 2021 (BP).

### VI gene clusters are associated with specific LUAD histopathology features in stRNA-seq data

Although the genes differentially expressed in tumors with VI were enriched for biological processes implicated in tumor intravasation, the RNA for sequencing was isolated from histologic sections selected to be representative of the predominant histologic pattern and were not selected to include invaded blood vessel(s). Given that VI foci represent such a small percentage of tumor volume (diameter ≤ 1 mm), we hypothesized that the VI signature we identified reflected molecular changes extending beyond the site of intravasation. To assess the spatial architecture of the expression of the VI signature and its association with LUAD histopathology features we profiled 15 (post-QC) resected stage I and stage II LUAD samples (n=8 VI^-^, n=7 VI^+^) from 13 patient tumors by high-resolution stRNA-seq using the 10X Genomics Visium platform, including 3 sections containing invaded vessels directly in the 6.5 mm^2^ Visium capture area (Table S2). Of the VI^-^ tumors, 7/8 were NST. All six non-mucinous LUAD growth patterns (solid, cribriform, micropapillary, acinar, papillary, lepidic) were represented across the stRNA-seq capture areas along with three invasion types (VI, VPI, and STAS) (Fig. 2a). Adjacent normal appearing lung was present in 12/16 capture areas. The histopathology associated with VI^+^ tumors in the stRNA-seq data was generally representative of trends observed in our larger clinical cohort, although the micropapillary pattern tended to be underrepresented in VI^+^ tumors (p.adj=8.62×10^-14^) and the papillary pattern was overrepresented (p.adj=1.01×10^-11^) (Extended Data Fig. 4a). A solid growth pattern and the presence of desmoplastic stroma were both associated with angioinvasion (p.adj=6.38×10^-31^; p.adj=2.26×10^-18^). While VI co-occurred with other invasion types frequently in our clinical cohorts, we did not find more than one invasion type per capture area in the stRNA-seq dataset. In summary, our stRNA-seq dataset captures many of the pathology features that are known to occur in stage I LUAD and co-occur with VI.

**Figure 2.**
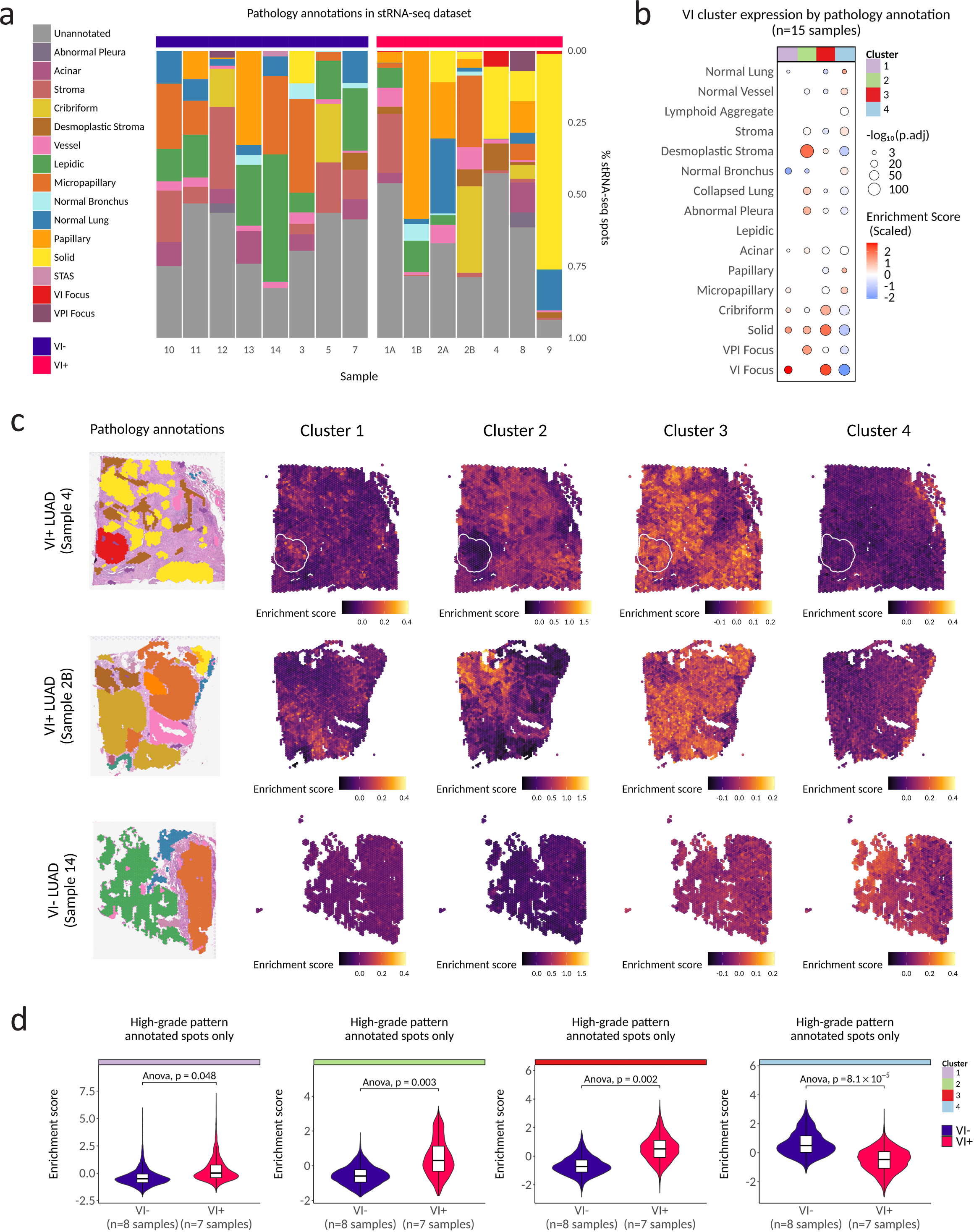
Spatial transcriptomics of early-stage LUAD reveals an association of VI gene clusters with specific LUAD histopathology features. **a**. Pathology annotations present across spatial transcriptomics (stRNA-seq) capture areas that passed QC (n=15). **b**. Association between expression of VI-associated clusters and pathology across all 15 samples. Only results with Bonferroni adjusted p values (p.adj) < 0.01 are shown. P.adj values were derived from a generalized binomial linear mixed effect model of VI cluster enrichment score as a function of spot pathology annotation with sample as a random effect. For visualization purposes the plot shows the mean of per-spot enrichment scores within a given region annotated as the same pathology. **c**. Pathology annotations (legend in a) in stRNA-seq samples and expression maps of VI gene expression clusters in representative samples from the VI^+^ (white outlines highlight VI foci) and VI^-^ LUAD. **d**. Spot-wise scoring of VI^-^ and VI^+^ stRNA-seq samples for expression of VI gene expression clusters in spots annotated with high-grade LUAD patterns (solid, micropapillary, and cribriform). P values are derived from a type II Anova of a linear mixed model predicting gene expression with sample as a random effect and high-grade pattern as a fixed effect.

After stRNA-seq quality control and filtering, we obtained expression estimates from 43,421 spots (50μm diameter) with a median depth of 8,548 unique molecular identifiers (UMIs) / spot (Extended Data Fig. 4b) that detected a median 3,996 genes / spot (Extended Data Fig. 4c). The low depth of sample 6 was likely due to tissue detachment that occurred during the stRNA-seq workflow, and this sample was removed prior to downstream analysis. Strong concordance was observed between pseudo-bulked stRNA-seq gene expression counts and bulk RNA-Seq counts of matched tumor tissues (spearman R=0.85) (Extended Data Fig. 4d). We next scored individual spots to create an enrichment score representing the activity of the genes from each of the four clusters in our bulk VI signature. Across all capture areas, spatially weighted correlation analysis of enrichment scores revealed that individual VI clusters were spatially distinct from one another (Extended Data Fig. 4e). These data suggest the distinct patterns of gene co-expression represented by the four clusters are also spatially distinct indicating that the biological processes that these gene expression patterns represent are spatially variable and are unlikely to represent redundant biology.

To investigate if the VI gene clusters are more strongly expressed in regions with different histopathologic feature annotations in the stRNA-seq data, we modeled the enrichment score of each cluster as a function of histopathologic annotation using a linear mixed model with the section as a random effect. Unexpectedly, despite the VI gene clusters being derived in bulk RNA-seq, we observed strong increased expression directly in VI foci of both cluster 1 (p.adj=9.02×10^-28^) and cluster 3 (p.adj=8.92×10^-73^). Cluster 1 and cluster 3 were also increased in regions annotated as high-grade patterns (solid (p.adj=8.87×10^-15^; 7.90^-75^) micropapillary (p.adj=3.94×10^-07^; 1.40×10^-23^) and cribriform (p.adj=4.23×10^-08^; 2.39×10^-58^)) (Fig. 2b). Cluster 2 was most noticeably expressed in desmoplastic stroma (p.adj=1.22^-122^) with a greater than 2.95-fold higher mean enrichment score than in stroma (p.adj=1.87×10^-25^). Conversely, expression of cluster 4 was strongly decreased in VI foci (p.adj=5.79×10^-79^), followed by desmoplastic stroma (p.adj=6.53×10^-50^) and showed strong positive enrichment in adjacent normal-appearing lung (p.adj=2.57×10^-04^), stroma (p.adj=8.90×10^-12^), and papillary (1.68×10^-13^). When enrichment scores were visualized directly on tissue sections, the independent spatial patterning of each cluster was apparent (Figs. 2c). Given that cluster 1 and 3 were significantly enriched directly in invasive foci in addition to high-grade patterns, we were interested to see whether the expression of these genes was also higher in VI^+^ tumors independently of aggressive pattern. When we analyzed only spots annotated with high-grade LUAD patterns, we found that cluster 1 (p=0.047), 2 (p=0.004) and 3 (p=0.003) were all higher and cluster 4 (p=6.9 x 10^-5^) was lower in spots annotated with high-grade patterns belonging to VI^+^ tumors, suggesting that our VI signature is capturing an angioinvasive phenotype that is independent of aggressive pattern and the invasive focus (Figure 2d). In total, these findings suggest that gene expression clusters of angioinvasive LUAD identified by bulk RNA-seq are differentially expressed between distinct histopathologic features *in situ*, show increased expression in the same tumor patterns if it is a VI^+^ tumor, and are not solely expressed in invaded blood vessels.

### The VI signature is composed of both tumor-specific and tumor-microenvironment changes reflective of angioinvasion

Our supervised analysis above relied upon detailed pathologic annotation of the tissue containing Visium spots, but spots were only labeled if there was a clear consensus on morphology, especially for the histologic pattern. In some cases, tumor areas may not have been annotated if they did not clearly contain canonical features of one of the six histologic patterns^30^. Additionally, for poorly differentiated patterns such as solid, the annotation may include admixed stromal or immune cells. With the aim to further delineate the spatial biology of the VI clusters, we estimated the cell type composition of each stRNA-seq spot with CytoSPACE by using a published single-cell RNAseq atlas of 295,813 annotated cells from 124 stage I LUAD samples (primary tumor and normal adjacent lung) and segmenting nuclei per spot with StarDist^31–34^. Visualization of cell types within the stRNA-seq data showed estimated cell types localizing with expected pathology structures (e.g., B cells with lymphoid aggregates) (Extended Data Fig. 5a). When we modeled predicted cell type proportion per spot as a function of histopathologic annotation using a linear mixed model with the section as a random effect we observed a significant enrichment of predicted B cells, Plasma cells, and other immune cell types with lymphoid aggregates (Fig. 3a). Normal bronchus had a significantly higher proportion of predicted ciliated cells, as expected. Spots annotated as VI foci had significantly higher proportions of predicted tumor cells while desmoplastic stromal regions had significantly higher proportions of peribronchial fibroblasts. Peribronchial, but not adventitial or alveolar fibroblasts, showed a significantly higher enrichment for a published NSCLC myofibroblastic-cancer-associated fibroblast (CAF) signature^35^ (Extended Data Fig. 5b). We found a trend toward a higher proportion of peribronchial fibroblasts belonging to VI^+^ samples (Fig. 3b), consistent with the tissue remodeling biology of VI cluster 2 described above. VI^+^ samples also had a trend toward a higher proportion of plasma cells and B cells. To validate these findings, we deconvoluted the bulk RNA-seq discovery cohort data using the same atlas. Although the predicted plasma cell proportions were associated with pathologist-annotated plasma cell grade on case-matched H&E images (p=1.37×10^-8^) (Extended Data Fig. 5c), we did not observe an association with VI^+^ tumors (p.adj=1), suggesting that plasma cell infiltration may be overrepresented in the stRNA-seq data. In contrast, we found a significantly higher predicted proportion of peribronchial fibroblasts (p.adj=0.005) and macrophages (p.adj=0.03) in VI^+^ tumors (Fig. 3c). To examine the associations between the expression of the VI gene clusters and cell types, we first generated cell type specific gene signatures from the reference atlas (Extended Data Fig. 5d). We then used spatially weighted regression to identify correlation between cell type signatures and our VI gene clusters across the stRNA-seq data after filtering low abundance cell types. Unsupervised clustering of the average spatially weighted correlation across all tumors revealed co-localization of cluster expression with distinct cell types (Fig. 3d). As expected from our previous observation of enrichment in desmoplastic stroma, VI cluster 2 was most spatially correlated with peribronchial fibroblasts, as well as other stromal cell types including endothelial and smooth muscle cells. In contrast, VI clusters 1, 3, and 4 were all tightly spatially correlated with epithelial cells, suggesting the contribution of these gene clusters to the VI signature is dependent more on cell state than cell type. Collectively, these results show that the VI gene clusters capture biological changes associated with angioinvasion from both tumor cells and the tumor microenvironment.

**Figure 3.**
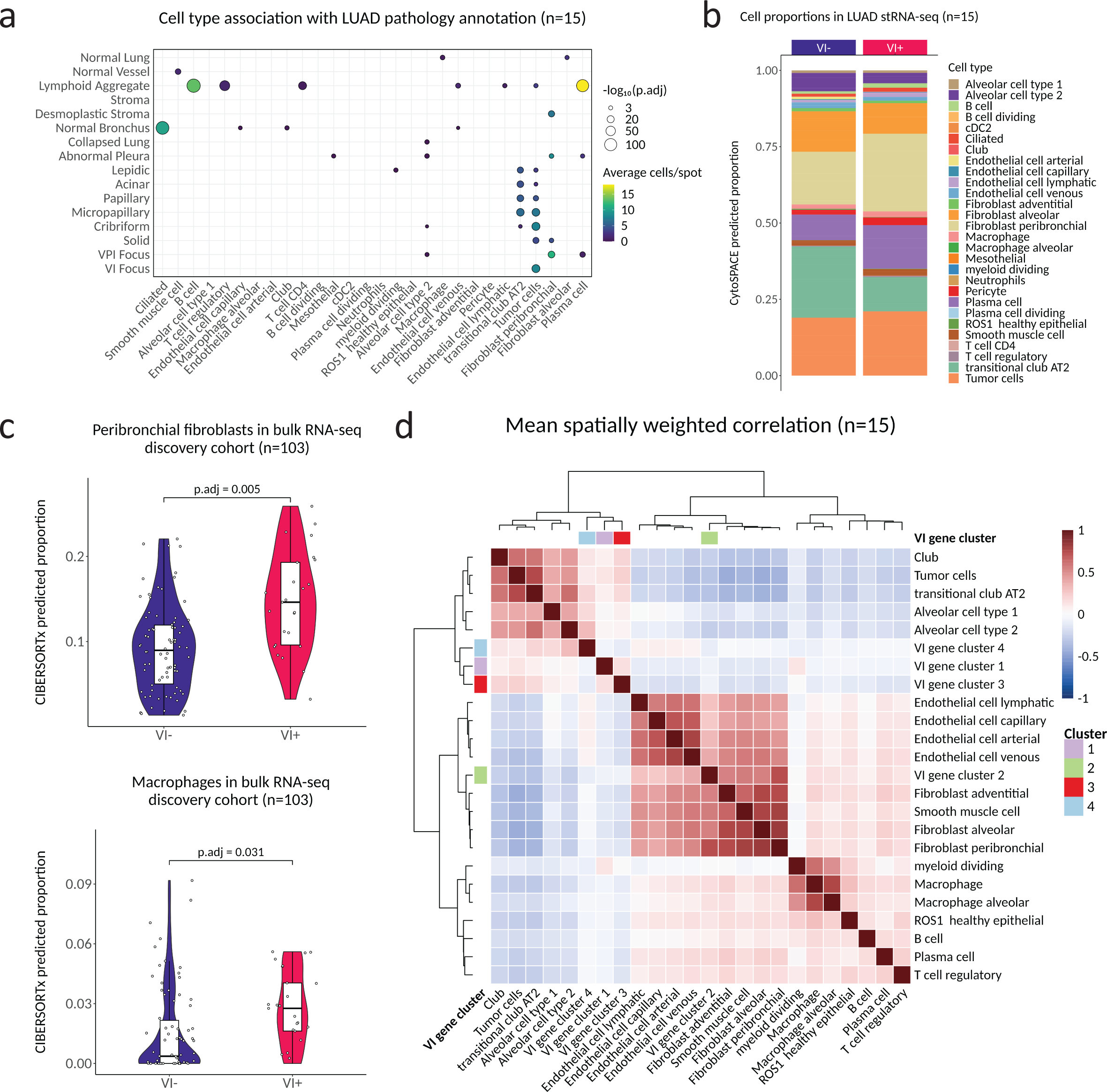
The VI signature is composed of both tumor-specific and tumor-microenvironment changes reflective of angioinvasion. **a**. Association between predicted stRNA-seq per-spot predicted cell type proportions and pathology across all 15 samples. Only results with p.adj < 0.01 are shown. P.adj values were derived from a linear-mixed model predicting cluster expression with sample as random effect. **b**. Cell type proportions by VI status in the stRNA-seq data, revealed by spot deconvolution. **c**. Peribronchial fibroblast and macrophage proportions by VI status in the bulk RNA-seq discovery cohort. **d**. Mean spatially weighted correlation of VI gene clusters and stage I LUAD cell type signatures from the Salcher et al. lung cancer atlas across all stRNA-seq samples.

### A VI predictor derived from the signature validates in an independent stage I LUAD cohort

Our finding of biologically and spatially distinct gene expression clusters associated with VI^+^ LUAD suggest that all four clusters provide orthogonal information and might be combined to form a predictor of angioinvasive Stage I LUAD. To test this hypothesis, we proceeded with a standard machine learning cross validation pipeline using only the samples from the discovery cohort (n=103; Fig. 4a). Training was performed in the discovery cohort (n=103) where the differentially expressed genes were first identified, using a cross-validation approach (Fig. 4a).

**Figure 4.**
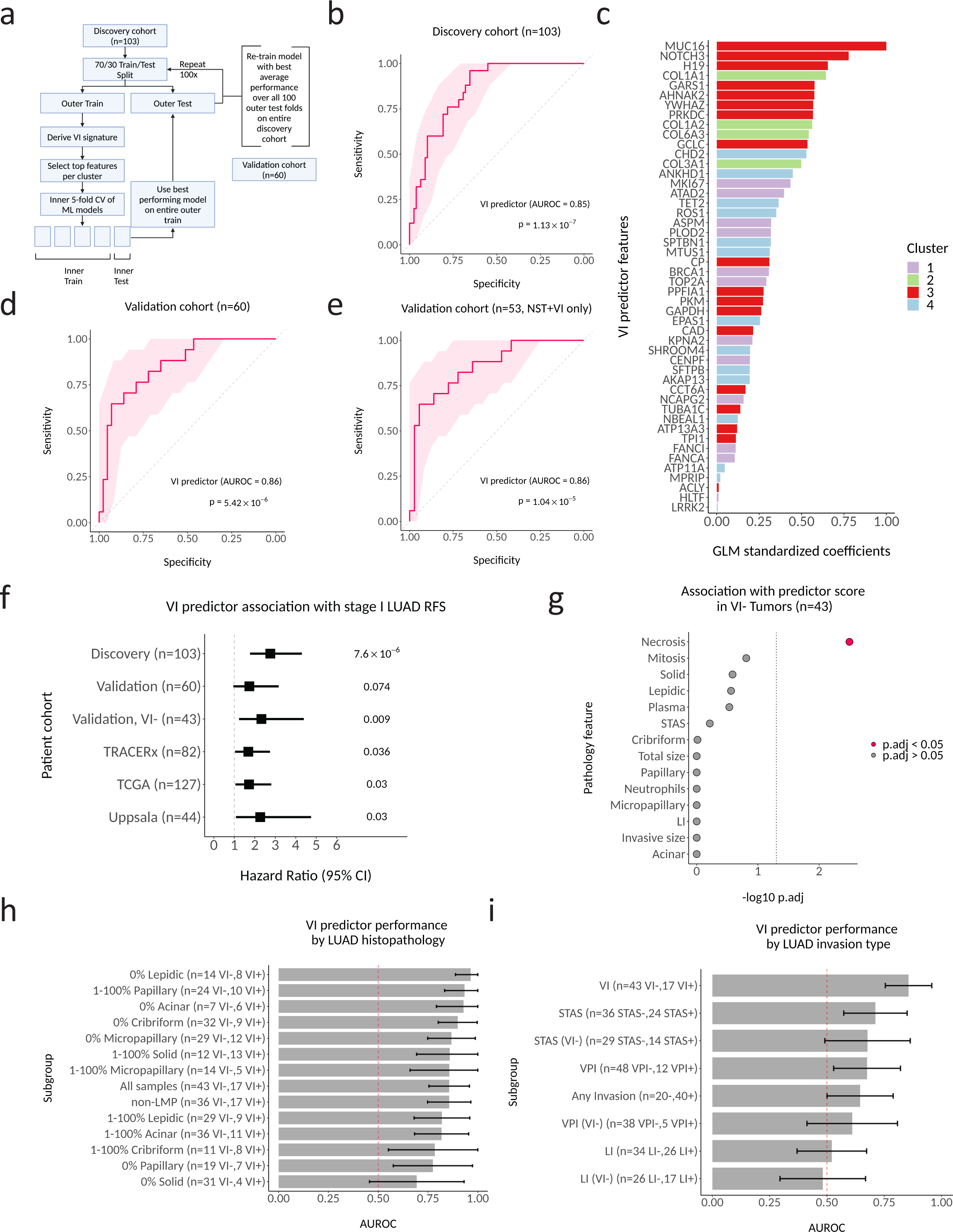
A VI predictor derived from the signature validates in an independent stage I LUAD cohort. **a**. Cross-validation approach for feature and model selection. **b**. Predictor performance for predicting VI in discovery cohort (n=103). The shaded region represents the 95% confidence interval (CI) of the sensitivity at different specificity points, computed with 2000 bootstrap replicates. P value obtained from Wilcoxon rank sum test. **c**. Feature importance for the final VI predictor. **d**. Predictor performance in the validation cohort (n=60). **e**. Predictor performance in the validation cohort in non-LMP tumors (n=53). **f**. Predictor association with RFS across stage I LUAD patient cohorts (discovery, validation, TRACERx, TCGA, Uppsala). P values were each calculated using univariate cox proportional hazards regression. **g**. Association of pathology features with VI predictor scores in VI-tumors within the validation cohort (n=43). A univariate linear model was used to calculate p.adj values for each pathology feature, with predictor scores as the dependent variable. **h**. Predictor performance in the validation cohort by histologic pattern (n varies). Error bar shows bootstrapped 95% CI. I. Predictor performance when classifying other LUAD invasion types in the validation cohort.

Predictors utilizing a binomial logit generalized linear model (GLM) with ridge regression performed the best on the internal cross-validation test sets within our discovery cohort (Extended Data Fig. 6a). When we trained a GLM with ridge regression on the full discovery cohort, a 48-gene predictor achieved an area under the receiver operating characteristic curve (AUROC) of 0.85 to separate VI^+^ and VI^-^ LUAD in the training set (Fig. 4b). When ranked by feature importance in the model, the top 3 genes in the predictor were from cluster 3 and included *MUC16*, *NOTCH3*, and *H19*, but only cluster 2 genes were significantly enriched among the top ranked genes overall (two-sided Kolmogorov-Smirnov test p = 0.02; Fig. 4C). The top features for clusters 1, 2, and 4 were *MKI67*, *COL1A1* and *CHD2*, respectively.

The predictor retained a similar performance in predicting VI^+^ LUAD when applied to our independent validation cohort (n=43 VI^-^, n=17 VI^+^) with an AUROC of 0.86 (Fig. 4d). It retained equivalent performance even after excluding LMP tumors (NST vs. VI only, AUROC 0.86) (Fig 4e). It was associated with histologic grade and 7-year recurrence free survival (HR=1.98, 95% CI 1.09–3.60, Cox regression p=0.024) (Extended Data Fig. 6b and Fig. 4f).

Additionally, it was also a significant predictor of RFS among VI^-^ LUAD tumors (HR=2.76, 95% CI 1.41– 5.41, Cox regression p=0.003). We found necrosis was associated with higher predictor scores in the VI^-^ tumors in a univariate linear model (p.adj = 0.003) (Fig. 4g). In our validation set, the VI predictor retained good performance across tumor subsets that varied by histological growth patterns, with the best performance in samples that contained no lepidic pattern (0% lepidic, n=22, mean AUROC 0.96) and the worst performance in samples that contained no solid pattern (0% solid, n=35, mean AUROC 0.69) (Fig. 4h). It also showed variable performance in predicting other invasion types, and notably was not significantly better than a random classifier at detecting LI (Fig. 4i), supporting our hypothesis that angioinvasion is a different molecular process than lymphatic invasion. Although we did not have separate VI annotations for published LUAD RNA-seq datasets, in TRACERx, a longitudinal study of NSCLC, we observed a difference in predictor scores between LVI^+^ and LVI^-^ LUAD (p=0.0014) (Extended Data Fig. 6c). Interestingly, VI predictor scores were higher in tumors from patients with detected preoperative ctDNA (p=5.9×10^-4^; Extended Data Fig. 6d). Finally, higher VI predictor scores were significantly associated with decreased 5-year RFS in TRACERx (Fig. 4f; HR=1.69, 95% CI 1.04–2.75, Cox regression p=0.036). Higher VI predictor scores additionally associated with decreased overall survival in stage IA LUAD tumors from both The Cancer Genome Atlas (TCGA) (n=127) (HR=1.72, 95% CI 1.05–2.81, Cox regression p=0.03) and Uppsala (n=44) (HR=2.11, 95% CI 1.05–4.27, Cox regression p=0.037) cohorts^36,37^.

### The VI predictor is robust to intra-tumor heterogeneity

Our stRNA-seq analysis of the expression of VI-associated genes implies that each VI-associated cluster may be spatially distinct. This suggests that a VI predictor combining genes from each cluster might overcome intra-tumor heterogeneity (ITH). Overcoming intra-tumor heterogeneity is crucial for molecular biomarkers that sample only a portion of the tumor volume, such as those measured on tissue available from biopsies^38^. This is particularly important for detecting VI due to the small size of invaded vessels. When we divided stRNA-seq spots from VI^+^ samples into distal VI^+^ (spots >1mm beyond the edge of invaded vessels) and proximal VI^+^ (spots < 1mm from and including invaded vessels) (Fig. 5a), we found a significant difference in the enrichment score between VI^-^ and distal VI^+^ tumors for the predictor genes (p=0.028) (Fig. 5b). This supports the hypothesis that the predictor may be able to detect VI away from the invasive focus. Next, we used RNA-seq from multi-region sampling of stage I LUAD (n=63) in TRACERx to evaluate the variability of the VI predictor to random tissue sampling in a larger cohort^29^. Although a label for VI was not available for the TRACERx tumors, we observed strong correlation between predictor scores from two randomly selected paired regions (n=126) across all tumors (R=0.87, p<2.2×10^-16^) (Fig. 5c). When we filtered the dataset to remove lowly expressed genes and ranked the remaining 15,800 genes in the dataset by correlation between paired regions, the 48 genes present in our predictor were significantly enriched among those with the least ITH (p=2.89×10^-6^) (Fig. 5d).

**Figure 5.**
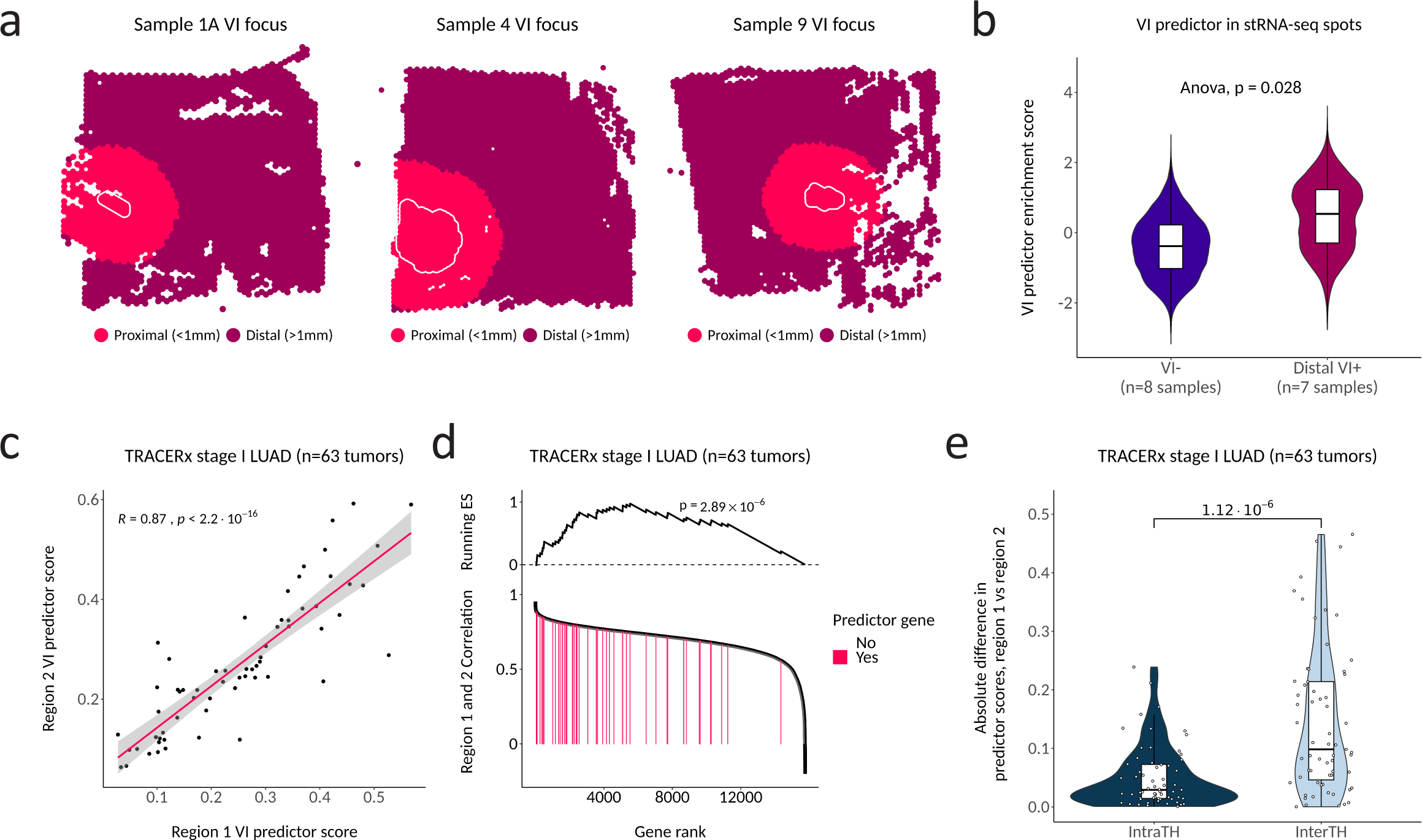
The VI predictor is robust to intra-tumor heterogeneity. **a**. Examples of binning VI^+^ tumors into distal VI^+^ (spots > 1mm outside invaded foci boundary) and proximal VI^+^ (spots < 1mm from and including the invaded foci). Any VI^+^ tumors that did not contain VI foci in the capture area were considered as proximal VI^+^. **b**. Spot-wise expression of the VI predictor (predictor up gene enrichment minus predictor down gene enrichment) between distal VI^+^ and VI^-^ tumors. P values are derived from a type II Anova of a linear mixed model predicting gene expression with sample as a random effect. **c**. Correlation between VI predictor scores from randomly sampled tumor-matched regions (n=126) of stage I LUAD cases from TRACERx, a study of NSCLC in which RNA-seq was performed on multiple tumor regions for a subset of tumors. P value shown for Spearman rank coefficient. **d**. The enrichment of the 48 VI predictor genes among all filtered genes ranked by correlation between two randomly selected regions of the same tumor. P value calculated by gene set enrichment analysis (GSEA). **e**. Differences in scores between unmatched regions (inter-tumor heterogeneity) and matched regions (intra-tumor heterogeneity). For inter-tumor heterogeneity, the absolute difference was calculated between two regions that were randomly sampled from two different tumors. This was repeated 63 times to make it equivalent to the intratumor heterogeneity calculation. P value calculated by Wilcoxon rank sum test.

Furthermore, inter-tumor heterogeneity of the VI predictor score (as measured by the absolute difference of scores from regions of two randomly selected tumors) was significantly higher than ITH (as measured by the absolute difference of scores from two randomly selected regions within the same tumor) (p=1.12×10^-6^) (Fig. 5e). Taken together, these results support our observations made from the stRNA-seq data that our VI signature represents a global shift in molecular aggressiveness toward an angioinvasive phenotype and that these changes can be leveraged to predict VI^+^ LUAD from any region of the tumor.

## DISCUSSION

Vascular invasion (VI) is a hallmark of tumor progression and a strong independent predictor of recurrence in stage I LUAD, suggesting that VI might result in considerable alterations in gene expression. To interrogate this, we used RNA-seq and stRNA-seq to profile new patient cohorts of stage I LUAD tumors that had undergone detailed pathologic assessments. We identified a gene expression signature of VI in stage I LUAD that is replicated across datasets, patient cohorts, and sequencing technologies. *In situ*, components of the VI gene signature can be localized to histopathologic structures outside of the invaded vessel. A VI predictor developed from this signature discriminates between VI^+^ and VI^-^ tumors in an independent validation cohort and is independent of histologic growth pattern. We also demonstrated that the VI predictor is robust to intratumor heterogeneity (ITH). This property has the potential to be exploited to develop a new preoperative biomarker for angioinvasive stage I LUAD.

Gene expression alterations associated with VI clustered into four distinct patterns of co-expression with unique biology. Gene clusters that had increased expression in VI^+^ tumors were enriched for cancer hallmarks including cell cycle (Cluster 1), EMT/angiogenesis (Cluster 2), and hypoxic tumor metabolism (Cluster 3). Tumor hypoxia is well documented as a tumor phenotype with poor prognosis across multiple cancer types and has been shown to be moderately associated with solid histopathology in LUAD^39,40^. Genes that were decreased in VI^+^ tumors (Cluster 4) included genes responsible for tumor suppression, control over cell-cell adhesion and immune responses. We also observed a greater number of transcriptional changes associated with VI than LI and aspects of these signatures had unique biology supporting previous reports^8,21^ that stage I LUAD VI and LI are histologic categories with distinct behavior. Specifically, expression of Cluster 2 which was enriched for genes involved in tissue remodeling and EMT and was elevated in VI^+^ tumors was not elevated in LI^+^ tumors. Lymphatics are thin walled with less surrounding connective tissue than blood vessels. The absence of elevated Cluster 2 expression in LI may reflect these microstructural differences that require less extensive tissue remodeling or may rely upon alternate mechanisms of invasion. Interestingly like VI, LI was associated with increased expression of genes in Cluster 1 and Cluster 3 and decreased expression of genes in Cluster 4. Despite a large spatial transcriptomics dataset containing many different histologic features, we were not able to profile an adequate area of LI. Future efforts may seek to separate additional biological signals or tumor subtypes that are associated with LUAD lymph node spread, given that tumor LI is not a reliable indicator of lymph node metastasis.

Because our bulk RNA-seq did not specifically profile dissected invaded blood vessels, it was not clear whether our VI signature was associated with invaded vessels directly or was a tumor-wide property of angioinvasion. Within the stRNA-seq data, expression of cluster 2 was most strongly enriched in regions of desmoplastic but not normal stroma, while expression of cluster 4 was enriched in low-grade histologic patterns and normal pulmonary structures. Desmoplastic stroma is a natural response of surrounding normal tissue to invasive tumor and involves widespread tissue remodeling^41^. The spatial expression pattern of Cluster 2 is consistent with the large number of genes involved in EMT and tissue remodeling in Cluster 2. The association between VI and desmoplastic stroma has been documented in thyroid carcinoma^42^, but we were not able to examine the co-occurrence in our LUAD cohort as the presence of desmoplastic stroma is not routinely assessed in LUAD. However, desmoplastic stroma has been associated with higher rates of distant metastasis or other prognostically unfavorable histologic features in other cohorts^43,44^.

We detected the expression of Cluster 1 and 3 genes within invaded blood vessels, but cluster 3 gene expression was also enriched in areas with solid and cribriform growth patterns more distal to the site of VI. Interestingly, within just the tumor regions exhibiting aggressive growth patterns, expression of Clusters 2 and 3 were significantly increased in VI^+^ tumors relative to regions of similar histology from VI^-^ tumors. Collectively, these data suggest that aspects of VI-associated gene expression occur distally to the site of VI and are not just a proxy for poorly differentiated tumors, but a signature of the most aggressive early-stage tumors. An inherent limitation of spot-based stRNA-seq data is the lack of single cell resolution, but spot deconvolution of stRNA-seq data using cell type signatures derived from human single-cell RNA seq datasets allowed us to interrogate VI-associated differences in predicted cell type abundance and the colocalization of VI-associated gene expression and cell-type specific signatures. Tumors with VI showed higher proportions of peribronchial or myofibroblastic-cancer-associated fibroblasts (CAF), which we validated in our bulk RNA-seq discovery cohort. The expression of Cluster 2 was most spatially correlated with CAFs and other stromal cell types while Cluster 1, 3, and 4 were spatially correlated with epithelial cell types.

We developed a VI predictor that strongly discriminated between VI^+^ and VI^-^ tumors in an independent validation cohort and was independent of tumor histologic pattern. The top weighted genes in this predictor belonged to Cluster 3 and included *MUC16*, *NOTCH3*, and *H19*. *MUC16*, which encodes CA125, enhances LUAD aggressiveness through downstream inactivation of *p53*^45^. Although serum CA125 is used as a diagnostic biomarker for patients at high risk of ovarian cancer, its association with lung cancer prognosis has not been demonstrated^46^. NOTCH pathway activity is associated with poor LUAD patient prognosis^47^ and *NOTCH1* is essential for LUAD tumor cell survival under hypoxia^48^. Depleting *NOTCH3* in LUAD also leads to reduced tumor cell invasion and significantly reduces the expression of *COL1A1*^49^, the top Cluster 2 gene in our VI predictor. The noncoding RNA *H19* locus is a tumor suppressor^50^. In lung cancer it is induced by the oncogene *Myc*, which drives tumorigenesis by binding near to the *H19* imprinting control region^51^. *H19* also represses *CDH1*, which is responsible for maintaining cell-cell adhesion^52^. The top Cluster 1 gene, *MKI67*, is expressed by proliferating cells and integral to carcinogenesis^53^. Finally, *CHD2*, the top Cluster 4 gene in our predictor, is a chromatin remodeler that along with *CHD1* and *CHD5* belongs to a family of tumor suppressor genes implicated in various cancers^54–56^, consistent with the decreased expression observed in VI^+^ LUAD. Collectively, many of the highly weighted genes in the VI predictor are implicated and may be accomplices in LUAD progression.

The VI predictor was also associated with recurrence-free survival (RFS) even in presumed VI^-^ tumors, where it was most associated with necrosis. Although necrosis can occur without VI, an *in vivo* model of breast cancer demonstrated that VI foci, necrosis, and circulating tumor cells are all temporally correlated and controlled by the tumor specific factor *Angptl7*, which promotes vascular permeability and remodeling^57^. Although this gene was not present in our VI predictor, the association in this study of predictor scores with VI foci, necrosis, and preoperative ctDNA status may suggest a similar temporal correlation in LUAD. Historically, prognostic gene signatures in lung cancer have suffered from poor reproducibility and lack of independence from known risk factors^18,58^. As an alternative to developing a molecular predictor of such a complex and potentially context-specific phenotype as outcome, our approach is to develop a predictor of a specific invasive feature that determines the histopathologic grade of stage I LUAD. Similar strategies have had substantial clinical utility, for example in guiding breast cancer management^59^.

Finally, our analysis of the multi-region sampling data from the longitudinal TRACERx cohort suggests that the VI predictor is robust to ITH. The ability to measure the VI predictor in 50 micron stRNA-seq tissue spots distal to invaded vessels suggests that the VI predictor might be insensitive to tissue composition and measurable in very small tissue specimens. We anticipate that molecular prediction of VI using tumor tissue from resected LUAD tumors may improve reproducibility of histopathological grading. Furthermore, an ability to predict VI from preoperative lung cancer biopsy tissue could have an even more substantial impact by helping to guide precision lung cancer surgery^60^. Although none of the cohorts we analyzed received neoadjuvant therapy, preoperative detection of VI^+^ tumors may in the future also provide the opportunity to also guide these treatments, which are increasingly being evaluated in earlier stages of resectable NSCLC^61^. Concordance of WHO grade between LUAD pre-surgical biopsies and resected specimens is poor^62,63^. Prospective studies should assess the agreement of VI estimation from biopsy and resected tumor material using both histopathology grading and molecular prediction. Integration of the VI predictor with other biomarker modalities such as radiomics and pathomics is likely to further improve risk stratification of stage I LUAD.

## METHODS

### Clinical cohorts

#### Discovery cohort

A discovery cohort consisting of 192 resected tumors from 192 patients with 8^th^ edition TNM stage 0/1 LUAD and not treated with neoadjuvant therapy were included in this study. Cases were from Boston Medical Center (BMC) and Lahey Hospital & Medical Center (LHMC). The BMC and LHMC Institutional Review Boards approved this study (BU/BMC IRB H-37859; Lahey Clinic IRB-518308). Relevant clinical information on all patients included can be found in Table S1. The 192 tumors represented a wide range of sample ages, so a small pilot batch of 12 samples was selected to determine which tumors might yield useful data (4/5 of the samples removed post-QC belonged to this pilot batch). We then selected an additional 96 samples that yielded at least 83 ng of library. RNA-seq was performed on these 108 tumors, with 103 passing quality control.

#### Validation cohort

An independent validation cohort consisting of 61 resected tumors from 60 patients with 8^th^ edition TNM stage I/II LUAD and not treated with neoadjuvant therapy were included in this study. All samples were selected to be stage IA/IB at time of collection except for one tumor that was upstaged to stage IIA under the 8^th^ TNM edition. Cases were from LHMC. RNA-seq was performed on 61 tumors, with all passing quality control. StRNA-seq was performed on a subset of these 61 tumors and included 16 samples taken from 14 tumors (13 stage I tumors and 1 stage II tumor). Relevant clinical information on all stRNA-seq samples included can be found in Table S2. Only 8^th^ edition TNM stage I LUAD from the validation cohort was included in the bulk RNA-seq (n=60) analysis. Relevant clinical information on all patients included can be found in Table S3.

### Pathology review

An experienced thoracic pathologist (E.B.) reviewed all pathology cases. Vascular invasion (VI) was defined as luminal invasion of a vein or muscular artery either within or adjacent to the tumor. Tumor proportions of lepidic, acinar, papillary, micropapillary, and solid patterns were assessed in 5% increments with distinction of simple tubular acinar from complex and cribriform acinar patterns. Adenocarcinoma in situ (AIS) was assigned to purely lepidic tumors ≤3 cm whereas minimally invasive adenocarcinoma (MIA) was diagnosed when non-lepidic foci measured ≤0.5 cm as per World Health Organization (WHO) criteria. WHO-2021 grade was defined as G1, lepidic predominant with <20% high-grade patterns; G2, acinar or papillary predominant with <20% high-grade patterns; and G3, ≥20% high-grade patterns (solid, micropapillary and/or complex glands)^64,65^. Low malignant potential adenocarcinoma (LMP) was assigned as previously described^24^. LMP tumors were non-mucinous adenocarcinoma measuring ≤3 cm in total size, with ≥15% lepidic growth, and without nonpredominant high-grade patterns (≥10% cribriform, ≥5% micropapillary, ≥5% solid), >1 mitosis per 2 mm^2^, vascular, lymphatic or visceral pleural invasion, STAS or necrosis. AIS/MIA and LMP were analyzed together due to their identical outcome (100% 10-year disease specific survival). One LMP recurred after wedge-resection with a positive surgical margin. The tumor recurred at the staple line and was treated with SBRT, resulting in prolonged survival (over 10 years) without further recurrence or metastasis. No special type (NST) designation was given for all other tumors not classified as VI or LMP. Stage assignments were retrospectively made using the 8th edition of the American Joint Committee on Cancer (AJCC).

### Statistical analysis

All statistical analysis was performed with R version 4.2.1. Tables were created with the tableone package. P values were corrected for multiple hypothesis testing where appropriate using either Bonferroni adjustment (abbreviated as p.adj) or false-discovery rate correction (abbreviated as FDR) depending on the default method of the R package used.

### RNA-Seq library preparation, sequencing, and data processing

Total RNA was extracted from FFPE tissue with the Qiagen AllPrep DNA/RNA Universal Kit and exome-targeted sequencing libraries were prepared with the Illumina TruSeq RNA Exome Library Prep Kit. Sample sequencing was performed on the Illumina NextSeq 500 to generate paired-end 50-nucleotide reads (discovery cohort) or Illumina HiSeq 2500 to generate paired-end 75-nucleotide reads (validation cohort). Basespace (Illumina) was used for demultiplexing and FASTQ file generation. A Nextflow v21.10.6 RNA-Seq pipeline aligned reads using hg38 and STAR^66^ v2.6.0c. Counts were calculated using RSEM^67^ v1.3.1 using Ensembl v108 annotation. Quality metrics were calculated with STAR and RSeQC^68^. EdgeR was used to compute normalized data (library sizes normalized using trimmed mean of M-values). Genes with count per million (cpm) > 1 in at least 10% of samples were retained for further analysis. Samples were excluded if the transcript integrity number (TIN)^69^ calculated by RSeQC was > 2 standard deviations from the mean TIN. Combat-seq^70^ was used to correct a batch effect between collection sites (BMC and LHMC) in the discovery cohort. It was also used to correct a batch effect between sequencing batches (batch 1 and 2) within the validation cohort.

### Derivation of VI signature and biological annotation

Genes whose expression is associated with the presence of VI were derived from 103 tumors in the discovery cohort using a negative binomial generalized linear model via edgeR^71^. Our previously published novel grading system (LMP, NST, VI)^8,16^ was the independent variable and we compared VI and LMP groups using glmLRT(). Differentially expressed genes were identified using a false-discovery rate (FDR) threshold of 0.01. Four gene expression clusters were identified using the Ward2 method of hierarchical clustering of genes and samples based on Euclidean distance^72^. The top ten biological processes and pathways enriched in each of the four VI clusters were identified using Enrichr^73^ with queries to the following databases: MSigDB Hallmark 2020 and GO Biological Process 2021. To compare VI and LI-associated gene expression, a VI signature was first rederived after adding LI as a covariate to the edgeR model described above, and differentially expressed genes were again identified using an FDR threshold of 0.01 when comparing VI and LMP groups. To identify genes associated with LI, contrasts were set to LI using makeContrasts() and differentially expressed genes identifying using an FDR threshold of 0.01. Genes associated with LI were ranked by -log_10_ (p value) * the sign of the logFC. The VI gene cluster enrichment against this ranked list was calculated using the GSEA() function from the clusterProfiler^74^ R package and p values were adjusted with p.adjust() for Bonferroni correction.

### Spatial transcriptomics library preparation, sequencing, and data processing

16 samples from 14 patient tumors within the validation cohort were selected for 10x Genomics Visium for FFPE spatial whole-transcriptome profiling. All samples were selected to be stage IA/IB at the time of collection except for one that was upstaged to stage IIA under the 8^th^ TNM edition (this sample was excluded from the bulk RNA-seq validation cohort when evaluating the performance of VI predictor scores). Samples were chosen based on 1) the presence of pathological features of interest (e.g., VI foci, representative LUAD histologic patterns) and 2) more than 50% of RNA fragments being greater than 200 nucleotides (DV200) after extraction with the AllPrep DNA/RNA Universal Kit (Qiagen) during bulk RNA-seq library preparation. New 5 μm tissue sections were cut by a microtome, placed in a 42°C RNase-free water bath and manually transferred onto 6.5 x 6.5 mm tissue capture areas on Visium Spatial Gene Expression Slides (PN-1000185, 10x Genomics). Tissues were deparaffinized, H&E stained, imaged with a Leica Aperio AT2, and de-crosslinked according to the manufacturer’s recommended protocol, with the following user modifications made to deparaffinization to prevent tissue detachment: 1) the 15 min incubation step during deparaffinization was removed, 2) after immersion in 96% ethanol, one 85%, one 70% and one 50% ethanol immersion for 3 min each was added. Probe hybridization was performed with the Visium Human Transcriptome Probe Kit (PN-1000364), followed by probe ligation, extension and elution for downstream qPCR cycle determination and 10x library construction with the Visium FFPE Reagent Kit (PN-1000362). Sequencing was done on an Illumina NextSeq 2000. The 10x SpaceRanger pipeline was used to demultiplex and generate FASTQ files. FASTQ files were input into the SpaceRanger FFPE count algorithm along with matching H&E .tiff images for alignment to the reference probe set and generation of count matrices for each Visium capture area.

### Spatial transcriptomics pathology annotations

H&E images of sections captured for stRNA-seq analysis were annotated for pathological features of interest by an experienced thoracic pathologist (EB) using Loupe Browser v6 (10x Genomics).

### Spatial transcriptomics data analysis

stRNA-seq data was processed using the Seurat R package, unless otherwise specified^75^. Sample 6 was removed due to tissue loss that occurred during wash steps. Low-quality spots and spots not covered by tissue were filtered if < 250 genes were detected per spot. For the global sample analysis, samples were merged and normalized using SCTransform and scaled^76^. For individual stRNA-seq samples, spots were log-normalized. Spots were scored with gene signatures using the AddModuleScore function from the Seurat R package. Spatial differences in VI cluster expression between pathology regions was assessed with a generalized binomial linear mixed effect model of VI cluster enrichment score as a function of spot pathology annotation with sample as a random effect. Pathology regions with < 200 spots annotated were removed prior to analysis and all regions were downsampled to 200 spots each. Normal lung was used as the reference factor level. In order to bin spots into distal VI^+^ and proximal VI^+^, image annotation screening on the invaded vessel pathology annotations was performed with the plotSurfaceIAS() function from the SPATA2^77^ R package with a distance and bin width of 1 mm.

### Spot deconvolution with scRNA-seq data

Nuclear segmentation of high-resolution H&E images of tissue used for stRNA-seq analysis was performed with the StarDist^33,34^ algorithm using default settings with a prob_thresh set to 0.3. Per-spot cell number estimates were generated within Squidpy^78^. Spot deconvolution was performed with the CytoSPACE^31^ algorithm using the per-spot cell numbers as input and the Salcher stage I LUAD atlas^32^ as the reference scRNA-seq dataset. The Salcher atlas was downloaded from https://luca.icbi.at/ and was filtered to stage I LUAD samples only. Spatial differences in cell type predictions between pathology regions was assessed with a generalized binomial linear mixed effect model of cell type proportion as a function of pathology annotation with sample as a random effect after averaging per-spot cell type proportions within a given region annotated as the same pathology. Pathology regions with < 200 spots annotated were removed prior to analysis and all regions were downsampled to 200 spots each. Normal lung was used as the reference factor level.

### Deconvolution of bulk RNA-seq data

Deconvolution of the bulk RNA-seq discovery cohort was performed with CIBERSORTx^79^ using the Salcher stage I LUAD atlas as input to the signature genes matrix generation step. The signature genes matrix was created by taking the average expression for each cell type per sample in the atlas.

### Spatially weighted correlation analysis

Spatial transcriptomic spots cannot be assumed to be spatially independent; therefore, spatially weighted regression analysis is used to calculate correlation between gene signatures. Unlike simple linear regression, which produces a single coefficient per variable, spatially weighted regression allows for coefficients to change locally with tissue coordinates. Data points closer to the coordinates at a given location within a set window (defined by the bandwidth) are given more weight within the model. This window can then slide over the whole tissue to calculate local correlation coefficients per-spot between variables. First, we used Seurat’s AddModuleScore to generate per-spot expression scores for our VI gene cluster signatures and cell type signatures from the Salcher atlas. Cell types that did not have at least one predicted cell by deconvolution across at least 20% of stRNA-seq capture areas were considered too sparse and were filtered out. Cell type expression signatures were generated for each of the remaining cell types annotated in the atlas using Seurat’s FindAllMarkers function with default settings. We selected the top 50 most significantly differentially expressed genes for each cell type. Spatially weighted regression was performed for each sample using the gwss function from the GWmodel^80^ package 2.3.1 with default settings and a bandwidth set to five. The mean Spearman’s rho of each correlation pair across all tissue spots was then taken for each sample and finally averaged across all samples.

### VI predictor derivation

A nested cross-validation pipeline (Fig. 4A) was used to evaluate performance of different machine learning models using only the discovery cohort samples and to prevent overfitting model hyperparameters. To accomplish this, samples from the discovery cohort were first divided into outer train and test sets using a 70/30 split 100 times.

#### Feature selection

Genes with VI-associated expression were derived within each train split as described above for the analysis of the entire discovery cohort: this included both selecting genes with FDR < 0.01 and dividing the genes into four clusters using hierarchical clustering. The size of the gene set used for generating a VI predictor in each fold of the cross-validation was set to 48. To automate feature selection and to ensure that diverse VI-associated gene expression patterns were evaluated, genes were selected within each fold of the cross-validation such that the proportion of genes from each of the four clusters was the same as the proportion of genes in that cluster among all the genes with FDR < 0.01 in that cross-validation fold. The gene set size of 48 was arbitrarily selected to allow for a 10-fold reduction of the size of the original signature in the full discovery cohort.

#### Prediction model selection

The outer train split was then divided into inner train and inner test folds using 5-fold cross-validation. This was to allow for evaluating predefined model hyperparameters (hyperparameter search) without learning information about the test data (the outer test fold) that would be used to inform final model selection. Model selection was based on comparing methods using this 5-fold cross-validation within the inner fold of the train split. This was done using the AutoML interface from the h2o.ai^81^ package v.3.40.0.4 and compared models including generalized linear model (GLM), gradient boosting machine (GBM), extreme gradient boosting (XGBoost) and distributed random forest (DRF). The best performing model and hyperparameter combination in the inner test fold was selected to develop a predictor on the outer train split and then applied to the held out outer test set. This was repeated over 100 cross-validation iterations using the discovery cohort. The final prediction model was selected as the model with the highest mean AUROC across all cross-validation iterations. This prediction model method and its associated hyperparameters, together with the feature selection step, were used with the gene expression data from the entire discovery cohort to develop a final 48-gene VI predictor using a binomial logit GLM utilizing ridge regression.

### VI predictor validation

Genes in the validation cohort were subsetted to those in the filtered discovery cohort. The mean and variance of each gene’s log-transformed cpm in the validation cohort was adjusted to match the discovery cohort using reference ComBat^82^ with the discovery cohort as the reference batch. VI predictor scores were generated using the final 48-gene predictor and performance was evaluated by AUROC. For evaluating the VI predictor scores by histologic subgroup, samples in the validation cohort were filtered based upon the percentage of annotated histopathologic growth patterns.

### Survival analysis

Univariate Cox proportional hazards regression was performed using the survival R package.

#### Additional datasets for survival analysis

Pre-processed RNA-Seq and matching clinical data were downloaded using a Genomic Data Commons (GDC) query for TCGA LUAD using the TCGAbiolinks^83^ R package and filtered to 127 stage IA samples with data on overall survival. Pre-processed RNA-Seq and matching clinical data from the Uppsala NSCLC cohort^37^ were downloaded using the Gene Expression Omnibus (GSE81089) and filtered to 44 stage IA LUAD samples with overall survival data. Because the novel histopathology grading system is based upon the 8^th^ edition of the TNM stage I samples and our predictor was derived in TNM 8^th^ edition stage I samples, we focused our analysis on this subset. Therefore, since tumor size was not available for either cohort, we excluded stage IB samples because these included tumors up to 5 cm in previous TNM editions and may have been recommended for adjuvant therapy. Genes in both datasets were subsetted to those in the filtered discovery cohort and the mean and variance of log-transformed cpm was adjusted to match the discovery cohort using reference ComBat with the discovery cohort as the reference batch.

### TRACERx intra-tumor heterogeneity analysis

Processed RNA-seq data and clinical annotations from the TRACERx NSCLC cohort of multi-region tumor sampling data were downloaded from https://doi.org/10.5281/zenodo.7683605 and https://doi.org/10.5281/zenodo.760338629. Samples were filtered to TNM 7^th^ edition stage I LUAD tumors ≤ 4 cm with data from at least two regions. Genes were subsetted to those in the filtered discovery cohort and the mean and variance of log-transformed cpm was adjusted to match the discovery data using reference ComBat with the discovery cohort as the reference batch. VI predictions were generated as described above. To determine intratumor heterogeneity in predictor scores, we calculated the absolute difference in VI predictor score between regions of the same tumor, when all tumors (n=63) were randomly downsampled to two regions each (n=126 regions total). For inter-tumor heterogeneity, the absolute difference was calculated between two regions that were randomly sampled from two different tumors. This was repeated 63 times to make the number of observations equivalent to the intratumor heterogeneity analysis. Spearman correlation between predictor scores from regions of the same tumor were also calculated. Predictor gene enrichment among all genes ranked by region-region correlation was determined using GSEA.

## Data availability

Data will be made available upon publication.

## ACKNOWLEDGEMENTS

This work was supported by R01CA275015, U01CA196408 and the Ellison Foundation. Figures 1A and 4A were created with BioRender.com.

## Contributions

D.S. processed raw sequencing data, performed all bioinformatic and statistical analyses, and wrote the manuscript. D.S. and L.S. performed spatial transcriptomics experiments and were supported by S.A.M. D.S., J.Z., K.R-C., E.J.B., J.B., and M.E.L. conceived the idea for and designed the study. E.J.B. performed the pathology review and assembled the cohorts and pathological data. T.S. led RNA isolation and was supported by K.R-C. H.L. and S.Z. performed RNA QC evaluation and library preparation and were supported by G.L. A.L. performed library preparation and Illumina sequencing and was supported by Y.O.A. J.B. and M.E.L. jointly supervised bioinformatic and statistical analyses and helped edit the manuscript.

## Corresponding author

Correspondence should be addressed to Marc Lenburg.

## Ethics declarations

### Competing Interests

The remaining authors declare no competing interests.

## Extended Data Figure Legends

**Extended Data Fig. 1. VI is the stage I LUAD invasion type most associated with recurrence. a**. Association of novel grading (n=192 tumors) and **b**. WHO grading (n=186 tumors) systems with 7-year RFS in the stage I LUAD discovery cohort. Mucinous tumors were excluded from WHO grading. P values were calculated by log-rank test. **c-f**. Association of VI, STAS, VPI, and LI pathology, respectively, with RFS. **g**. Co-occurrence of invasion types in the stage I LUAD clinical cohort. **h**. Association of VI with RFS (n=183 tumors) when controlling for common clinical variables, collection site (LHMC – Lahey Hospital & Medical Center, BMC – Boston Medical Center) and the other invasion types. Patients with missing values for any of the covariates were excluded. P values derived from multivariate Cox regression. **i**. Association between invasion types and LUAD growth patterns. Bonferroni adjusted p values (p.adj) were calculated by Wilcoxon test **j**. Proportion of VI- and VI+ cases that recurred at different locations. **k**. Recurrence site-specific sub-distribution HR for VI (n=185 tumors). Patients with unknown recurrence site were excluded from the analysis. HRs were adjusted for gender, age, pack years, surgical procedure, and collection site. Vertical lines designate the 95% CI. P values were calculated by multivariable Fine-Gray regression.

**Extended Data Fig. 2. The four VI gene expression clusters individually predict VI in the discovery cohort even in the absence of LMP tumors. a**. ROC curves for predicting VI^+^ vs. VI^-^ tumors using the mean z-score of each gene expression cluster in the discovery cohort (n=103). **b**. ROC curves for predicting VI vs. NST tumors using the mean z-score of each gene expression cluster in the discovery cohort (n=80). LMP tumors were excluded from the analysis. P values are reported for the Wilcoxon test where the null hypothesis is that the AUROC is equal to 0.5.

**Extended Data Fig. 3. VI but not LI is associated with expression of tissue remodeling genes. a**. Co-expression heatmap of 133 genes differentially expressed between VI and LMP (FDR < 0.01) in a discovery cohort of stage I LUAD tumors (n=103), with LI as a covariate. Genes are annotated by the clusters defined in Fig 1b. Heatmap units are log counts per million (CPM) scaled by transcript. VI, vascular invasion; NST, no special type; LMP, low malignant potential; LI, lymphatic invasion. **b**. Heatmap of 6 genes associated with LI contrasts (FDR < 0.01). Genes are annotated if they belong to one of the clusters defined in Fig 1B. **c-f**. Gene-set enrichment analysis (GSEA) results of the 4 VI gene clusters against a ranked list of genes ordered by association with LI. P values were calculated by GSEA.

**Extended Data Fig. 4. StRNA-seq quality control and concordance with bulk RNA-seq. a**. The association of spot-wise pathology annotations with tumor-level VI status in stRNA-seq samples that passed QC (n=15) (red = over-represented; blue = under-represented). P.adj values were calculated by chi-square test. **b**. Features per spot and **c**. UMI counts per spot by sample in the stRNA-seq data prior to QC (n=16). Sample 6 failed QC and was excluded from downstream analysis. **d**. Correlation of mean gene expression between pseudo bulked stRNA-seq data and tumor-matched bulk RNA-seq data (n=15 matched samples). P value shown for Spearman rank coefficient. **e**. Mean spatially weighted correlation of spot-wise VI cluster enrichment scores across all stRNA-seq samples (n=15).

**Extended Data Fig. 5. Cell type deconvolution of stRNA-seq and bulk RNA-seq. a**. Representative images showing the association of cell type enrichment signatures with pathology. **b**. Sample-level enrichment scores (averaged over single cells) of NSCLC myofibroblastic-cancer associated, alveolar, and adventitial fibroblast signatures derived from Hanley et al. 2023 in fibroblast subpopulations from Salcher lung cancer atlas stage I LUAD samples. **c**. Proportions of plasma cells from deconvolution of the bulk RNA-seq discovery cohort, stratified by pathologist annotated plasma cell grade (n=102 with annotations). **d**. Correlation of cell type markers (using the average expression of the top 50 differentially expressed marker genes) in the stage I LUAD samples from the Salcher lung cancer atlas.

**Extended Data Fig. 6. VI predictor development and validation. a**. Cross-validation performance for the four AutoML models in the outer test fold. Dr1, distributed random forest; gbm, gradient boosting machine; glm, generalized linear model; xgboost, extreme gradient boosting. Error bars represent standard deviation (SD) of AUROCs across all cross-validation folds (n=100). **b**. VI predictor association with novel and World Health Organization (WHO) 2015 grading systems in the validation cohort (n=60 tumors). For the WHO analysis, mucinous tumors (n=2) were removed from grade 3 as this grading system excludes mucinous tumors. P values were calculated by Wilcoxon test. **c**. Association of predictor scores with LVI^+^ tumors and **d**. patients’ preoperative ctDNA status in stage I LUAD RNA-seq data from the TRACERx cohort.

## Supplementary Tables

**Table S1.**
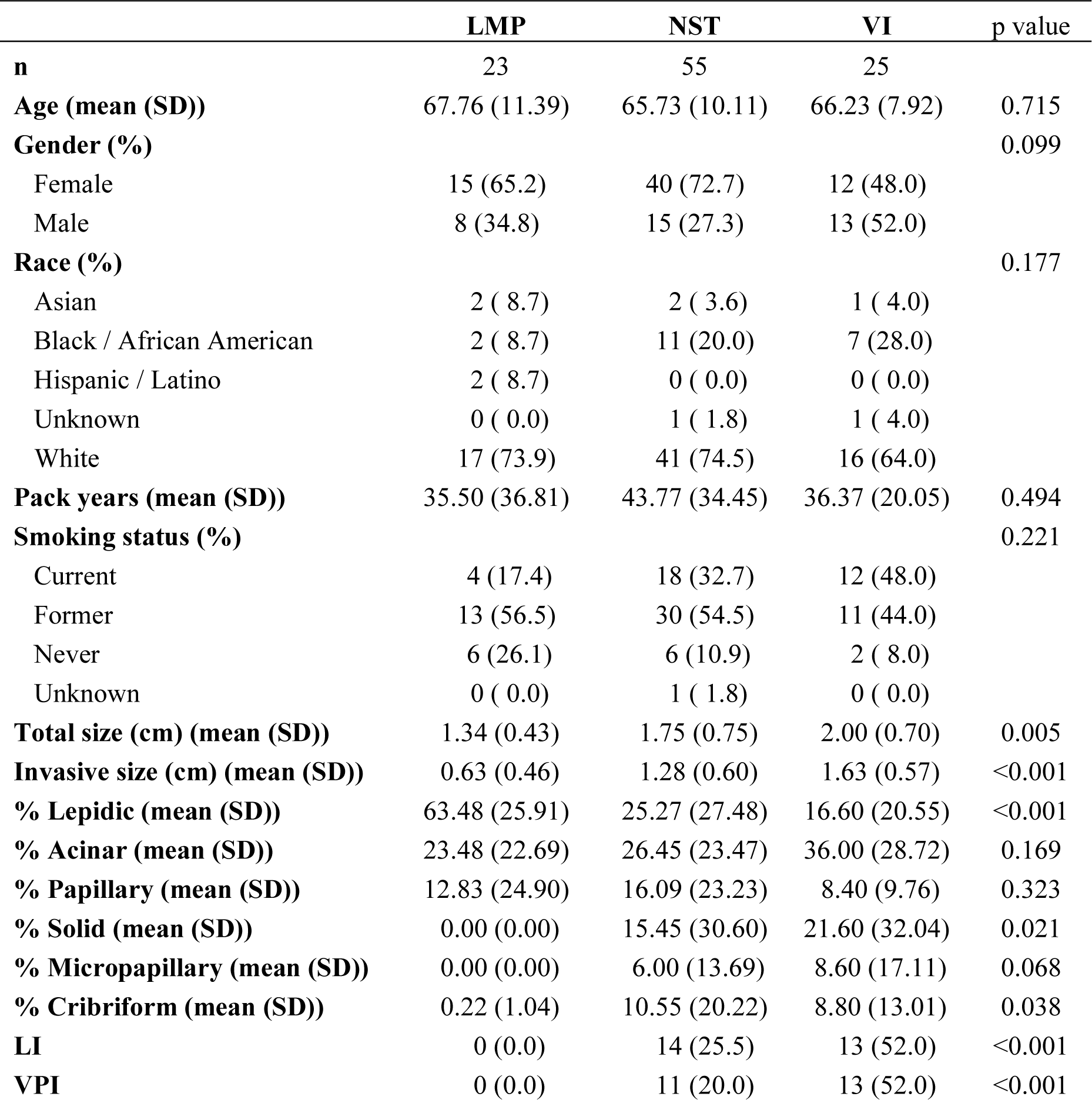
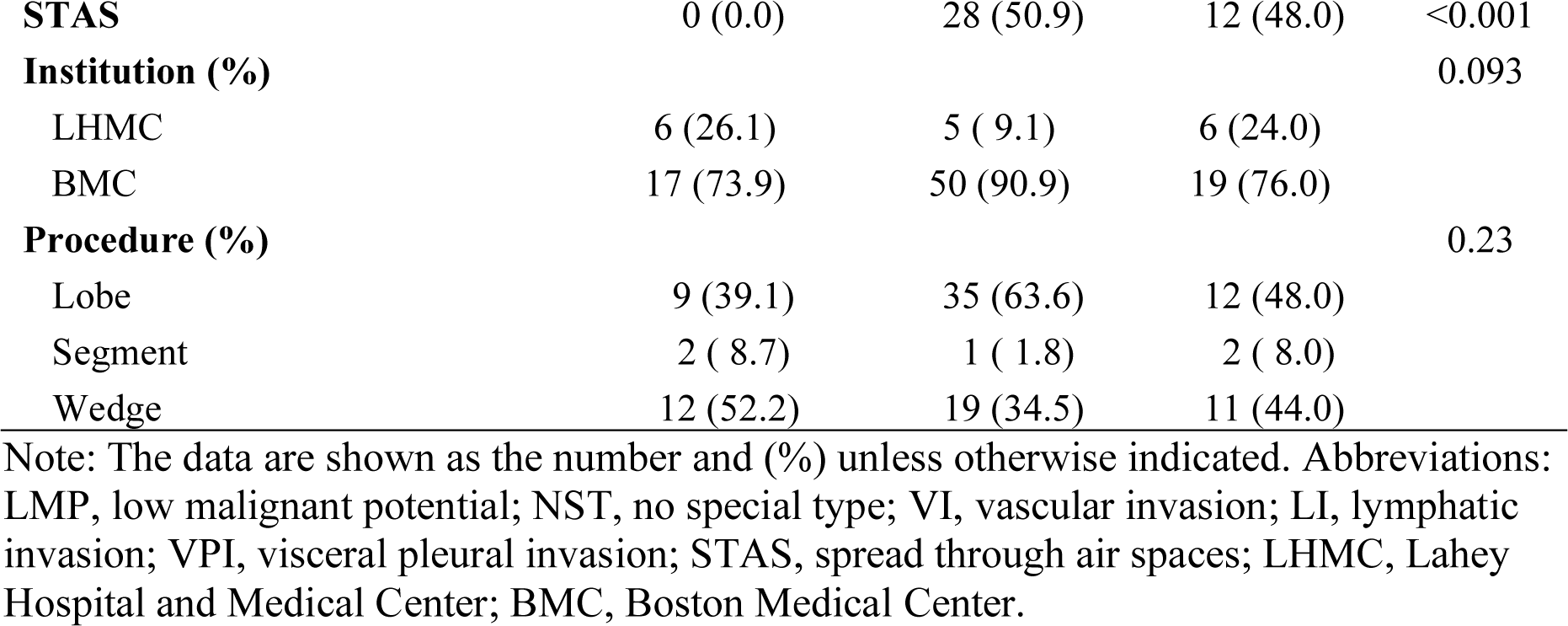
Clinical characteristics of resected stage I LUAD tumors with RNA-seq (post-QC) from the discovery cohort (n=103 tumors) using the novel histopathology classification.

|  | **LMP** | **NST** | **VI** | **p value** |
| --- | --- | --- | --- | --- |
| **n** | 23 | 55 | 25 |  |
| **Age (mean (SD))** | 67.76 (11.39) | 65.73 (10.11) | 66.23 (7.92) | 0.715 |
| **Gender (%)** |  |  |  | 0.099 |
| Female | 15 (65.2) | 40 (72.7) | 12 (48.0) |  |
| Male | 8 (34.8) | 15 (27.3) | 13 (52.0) |  |
| **Race (%)** |  |  |  | 0.177 |
| Asian | 2 ( 8.7) | 2 ( 3.6) | 1 ( 4.0) |  |
| Black / African American | 2 ( 8.7) | 11 (20.0) | 7 (28.0) |  |
| Hispanic / Latino | 2 ( 8.7) | 0 ( 0.0) | 0 ( 0.0) |  |
| Unknown | 0 ( 0.0) | 1 ( 1.8) | 1 ( 4.0) |  |
| White | 17 (73.9) | 41 (74.5) | 16 (64.0) |  |
| **Pack years (mean (SD))** | 35.50 (36.81) | 43.77 (34.45) | 36.37 (20.05) | 0.494 |
| **Smoking status (%)** |  |  |  | 0.221 |
| Current | 4 (17.4) | 18 (32.7) | 12 (48.0) |  |
| Former | 13 (56.5) | 30 (54.5) | 11 (44.0) |  |
| Never | 6 (26.1) | 6 (10.9) | 2 ( 8.0) |  |
| Unknown | 0 ( 0.0) | 1 ( 1.8) | 0 ( 0.0) |  |
| **Total size (cm) (mean (SD))** | 1.34 (0.43) | 1.75 (0.75) | 2.00 (0.70) | 0.005 |
| **Invasive size (cm) (mean (SD))** | 0.63 (0.46) | 1.28 (0.60) | 1.63 (0.57) | <0.001 |
| **% Lepidic (mean (SD))** | 63.48 (25.91) | 25.27 (27.48) | 16.60 (20.55) | <0.001 |
| **% Acinar (mean (SD))** | 23.48 (22.69) | 26.45 (23.47) | 36.00 (28.72) | 0.169 |
| **% Papillary (mean (SD))** | 12.83 (24.90) | 16.09 (23.23) | 8.40 (9.76) | 0.323 |
| **% Solid (mean (SD))** | 0.00 (0.00) | 15.45 (30.60) | 21.60 (32.04) | 0.021 |
| **% Micropapillary (mean (SD))** | 0.00 (0.00) | 6.00 (13.69) | 8.60 (17.11) | 0.068 |
| **% Cribriform (mean (SD))** | 0.22 (1.04) | 10.55 (20.22) | 8.80 (13.01) | 0.038 |
| **LI** | 0 (0.0) | 14 (25.5) | 13 (52.0) | <0.001 |
| **VPI** | 0 (0.0) | 11 (20.0) | 13 (52.0) | <0.001 |
| **STAS** | 0 (0.0) | 28 (50.9) | 12 (48.0) | <0.001 |
| **Institution (%)** |  |  |  | 0.093 |
| LHMC | 6 (26.1) | 5 ( 9.1) | 6 (24.0) |  |
| BMC | 17 (73.9) | 50 (90.9) | 19 (76.0) |  |
| **Procedure (%)** |  |  |  | 0.23 |
| Lobe | 9 (39.1) | 35 (63.6) | 12 (48.0) |  |
| Segment | 2 ( 8.7) | 1 ( 1.8) | 2 ( 8.0) |  |
| Wedge | 12 (52.2) | 19 (34.5) | 11 (44.0) |  |

Note: The data are shown as the number and (%) unless otherwise indicated. Abbreviations: LMP, low malignant potential; NST, no special type; VI, vascular invasion; LI, lymphatic invasion; VPI, visceral pleural invasion; STAS, spread through air spaces; LHMC, Lahey Hospital and Medical Center; BMC, Boston Medical Center.

**Table S2.** Clinical characteristics of resected early-stage LUAD tumors with stRNA-seq (post-QC) from the discovery cohort (n=15 samples).

| Sample ID | Batch | 8th Stage | Novel Grade | Age | Sex | Smoking Status | RNA-seq | VI (Capture Frame) |
| --- | --- | --- | --- | --- | --- | --- | --- | --- |
| 1A | 1 | IB | VI | 68 | M | Former | Yes | 1 |
| 1B | 2 | IB | VI | 68 | M | Former | Yes | 0 |
| 2B | 2 | IA3 | VI | 68 | F | Current | Yes | 0 |
| 2A | 2 | IA3 | VI | 68 | F | Current | Yes | 0 |
| 3 | 2 | IA1 | NST | 71 | M | Former | Yes | 0 |
| 4 | 2 | IB | VI | 66 | M | Former | Yes | 1 |
| 5 | 2 | IA1 | NST | 72 | M | Former | Yes | 0 |
| 7 | 2 | IA1 | LMP | 52 | F | Current | Yes | 0 |
| 8 | 1 | IB | VI | 66 | M | Former | Yes | 0 |
| 9 | 2 | IA2 | VI | 63 | F | Current | Yes | 1 |
| 10 | 2 | IA2 | NST | 69 | M | Former | Yes | 0 |
| 11 | 1 | IIA | NST | 63 | F | Former | Yes | 0 |
| 12 | 2 | IB | NST | 65 | F | Former | Yes | 0 |
| 13 | 1 | IA1 | NST | 72 | F | Former | Yes | 0 |
| 14 | 2 | IA2 | NST | 75 | F | Former | Yes | 0 |

Note: Abbreviations: LMP, low malignant potential; NST, no special type; VI, vascular invasion; LI, lymphatic invasion; VPI, visceral pleural invasion; STAS, spread through air spaces; LHMC, Lahey Hospital and Medical Center; BMC, Boston Medical Center.

**Table S3.**
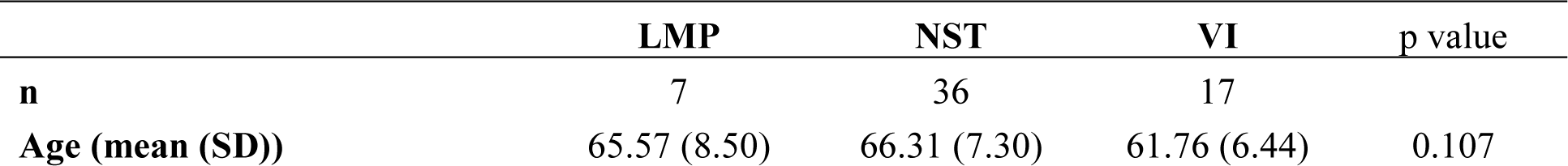
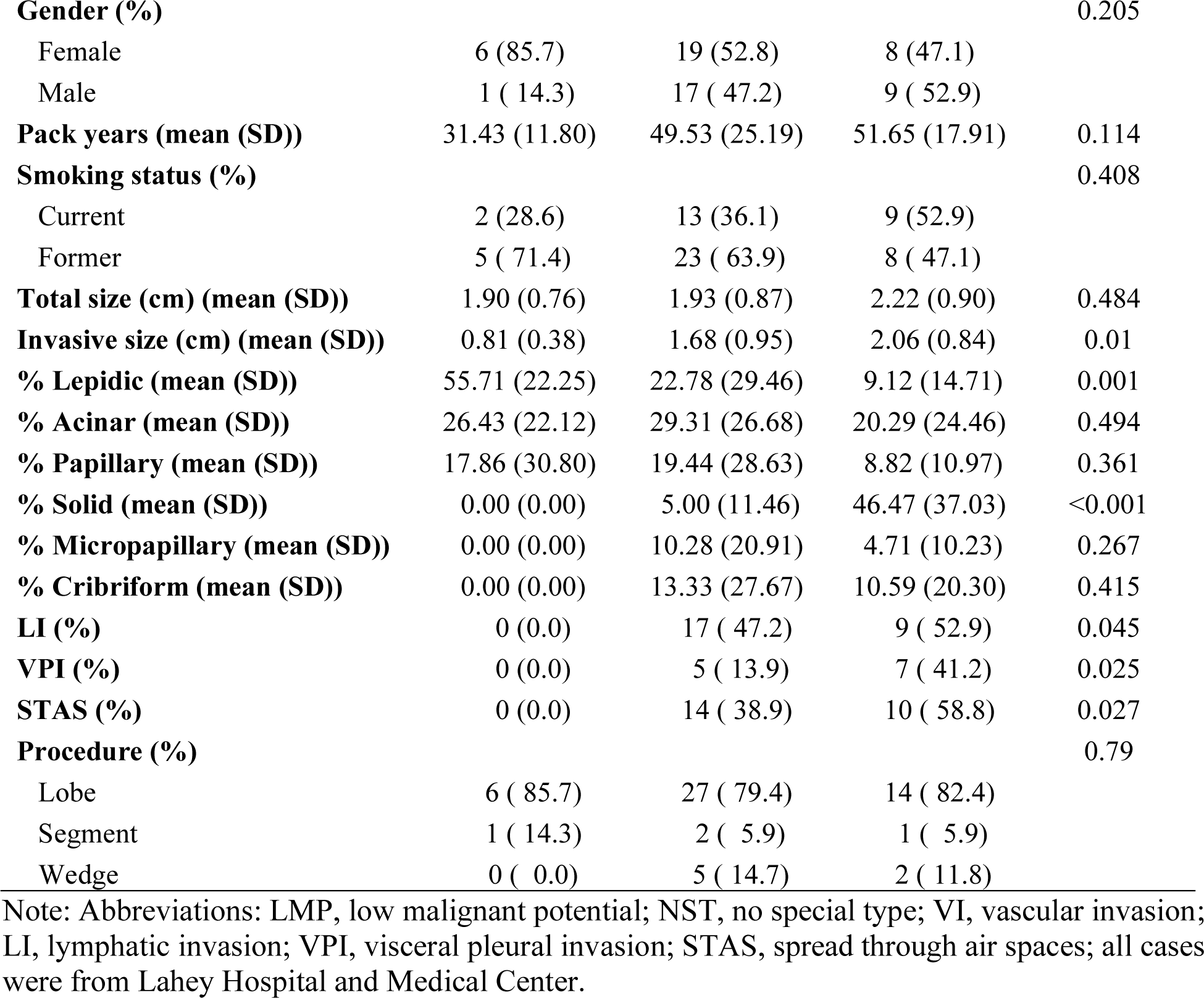
Clinical characteristics of resected stage I LUAD tumors with RNA-seq (post-QC) from the validation cohort (n=60 tumors) using the novel histopathology classification.

## Notes

### Competing Interest Statement

The authors have declared no competing interest.

