## Supplementary figures and images for "Identification of a gene expression signature of vascular invasion and recurrence in stage I lung adenocarcinoma via bulk and spatial transcriptomics"

### Extended Data Fig. 1

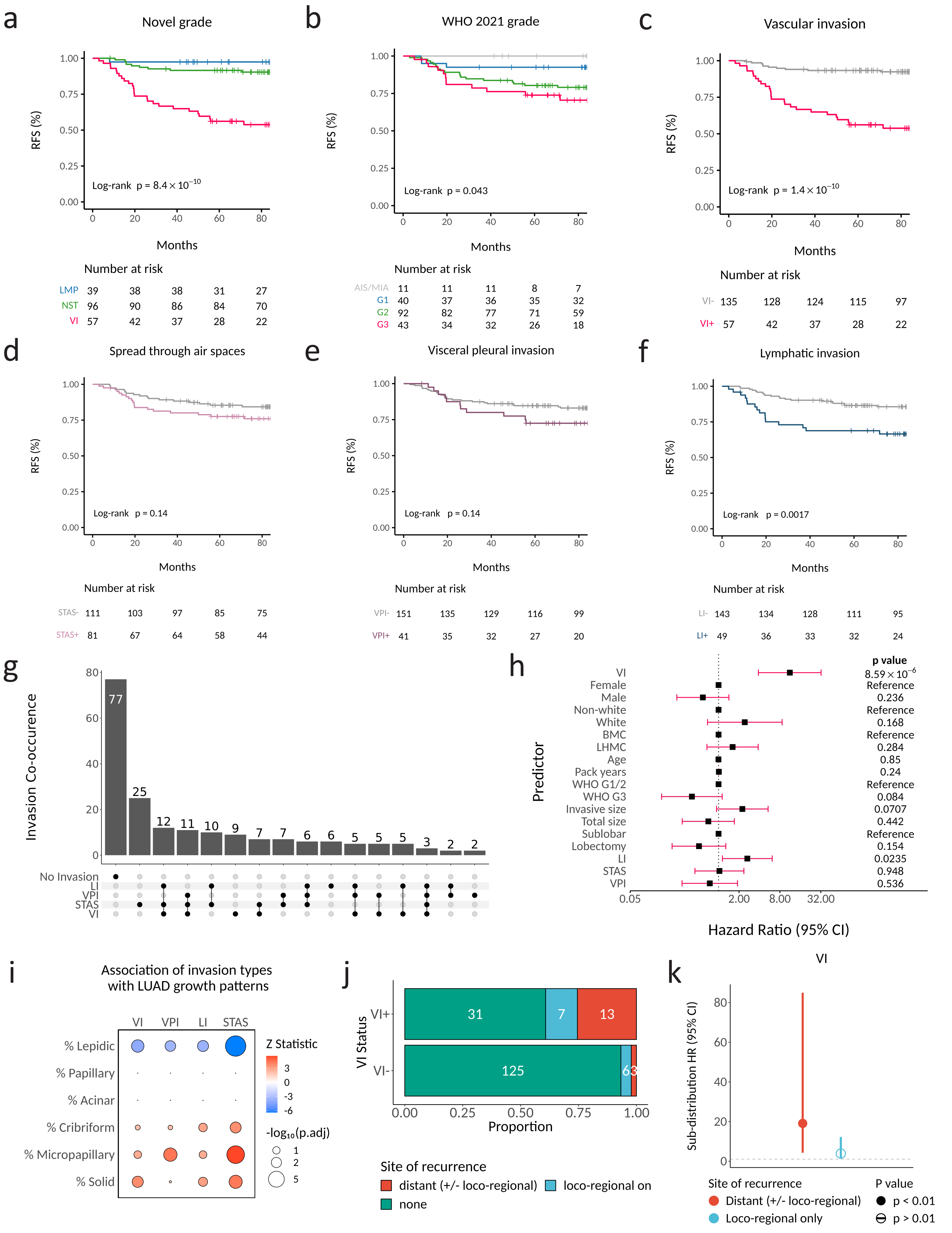


**Extended Data Fig. 1. VI is the stage I LUAD invasion type most associated with recurrence.**

### Extended Data Fig. 3

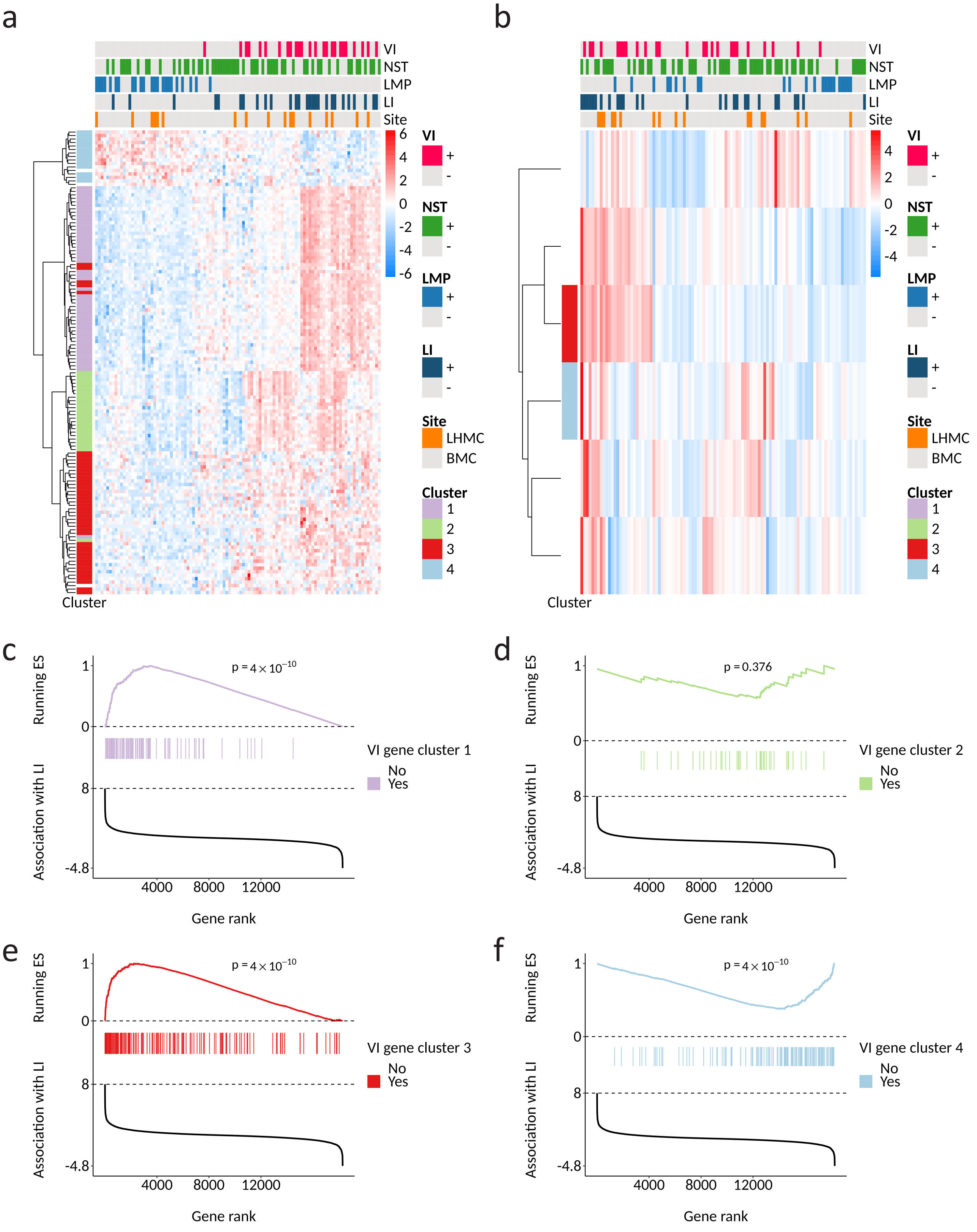


**Extended Data Fig. 3. VI but not LI is associated with expression of tissue remodeling genes.**

### Extended Data Fig. 4

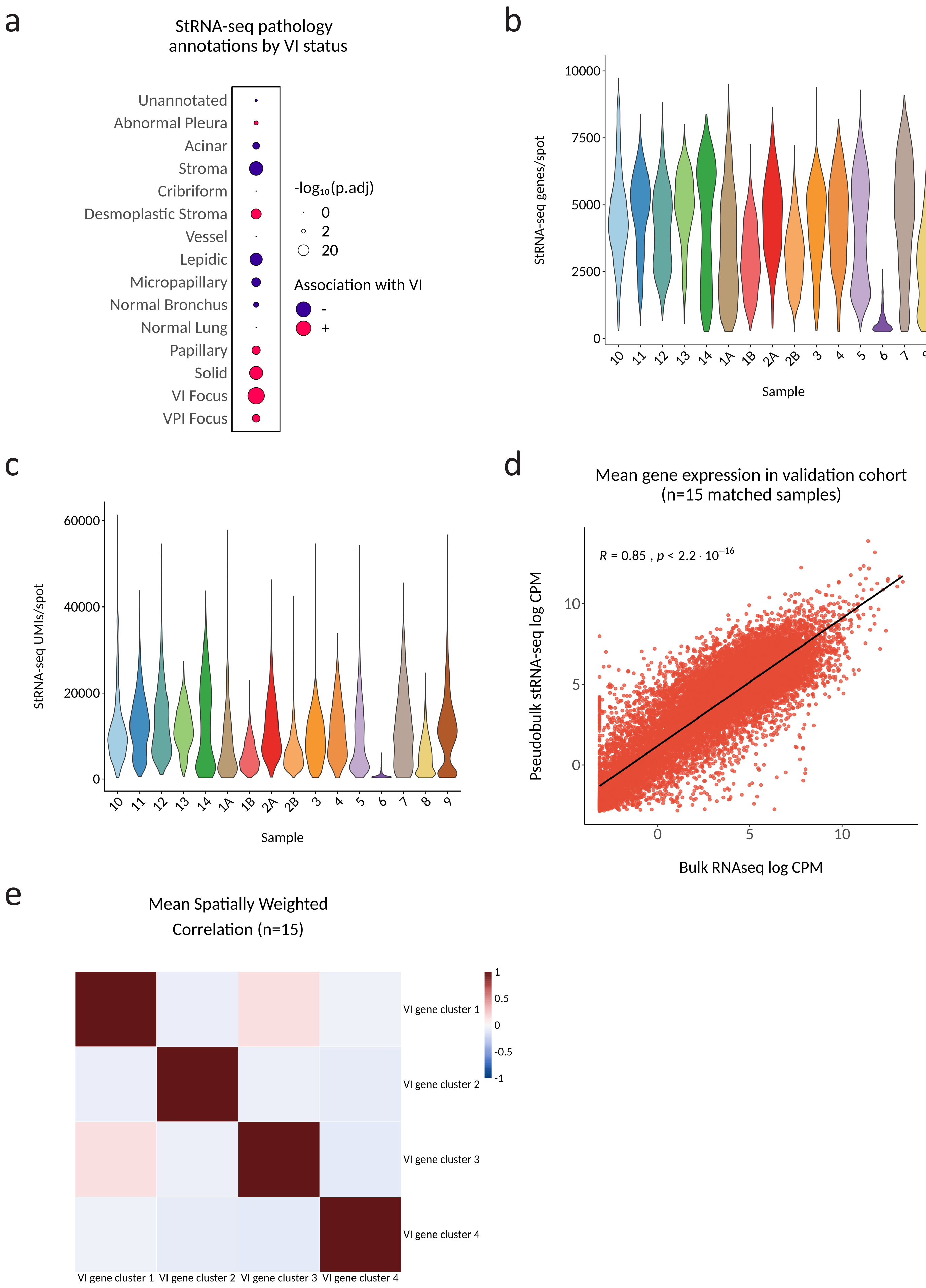


**Extended Data Fig. 4. StRNA-seq quality control and concordance with bulk RNA-seq.**

### Extended Data Fig. 5

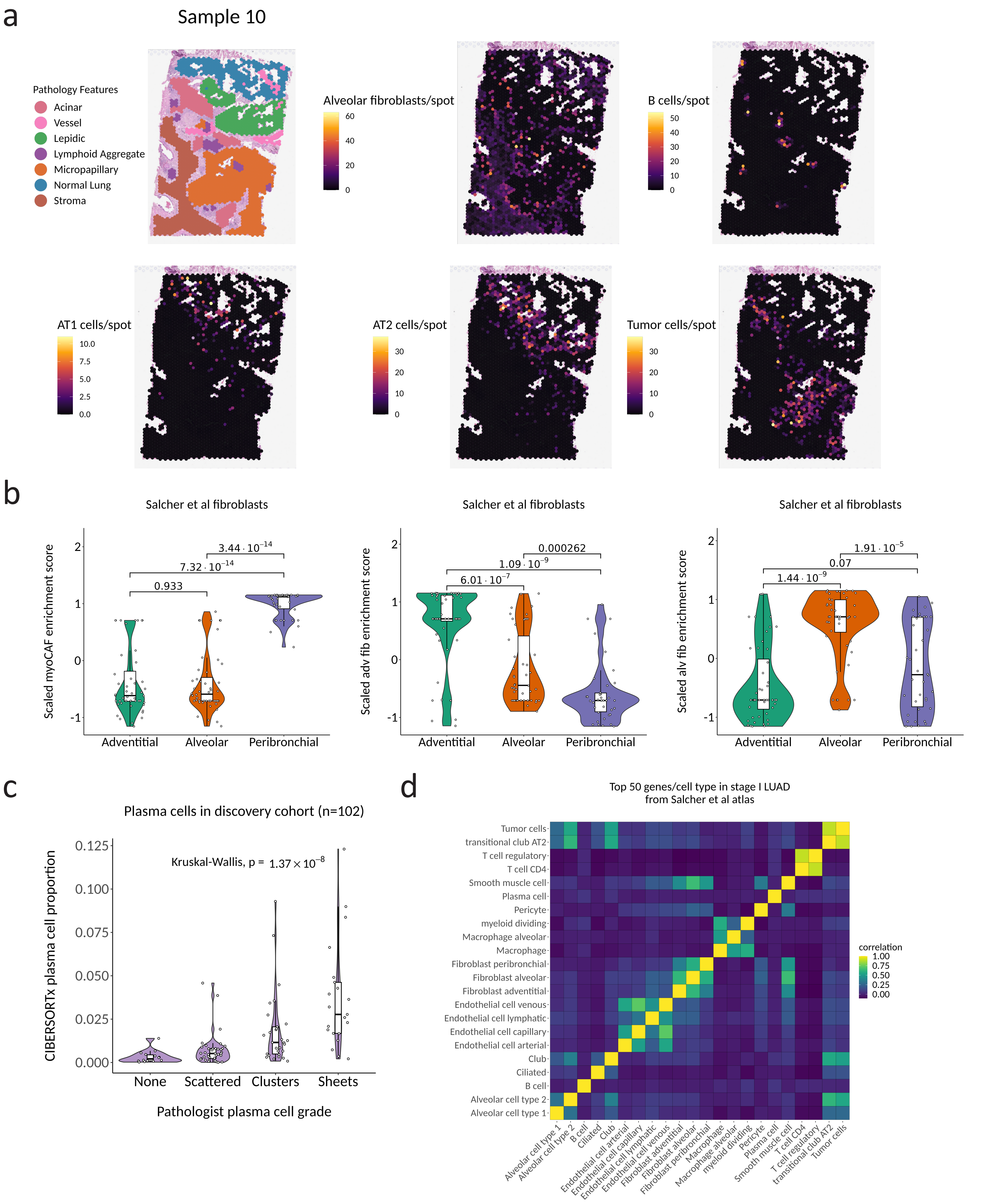


**Extended Data Fig. 5. Cell type deconvolution of stRNA-seq and bulk RNA-seq.**

### Extended Data Fig. 6

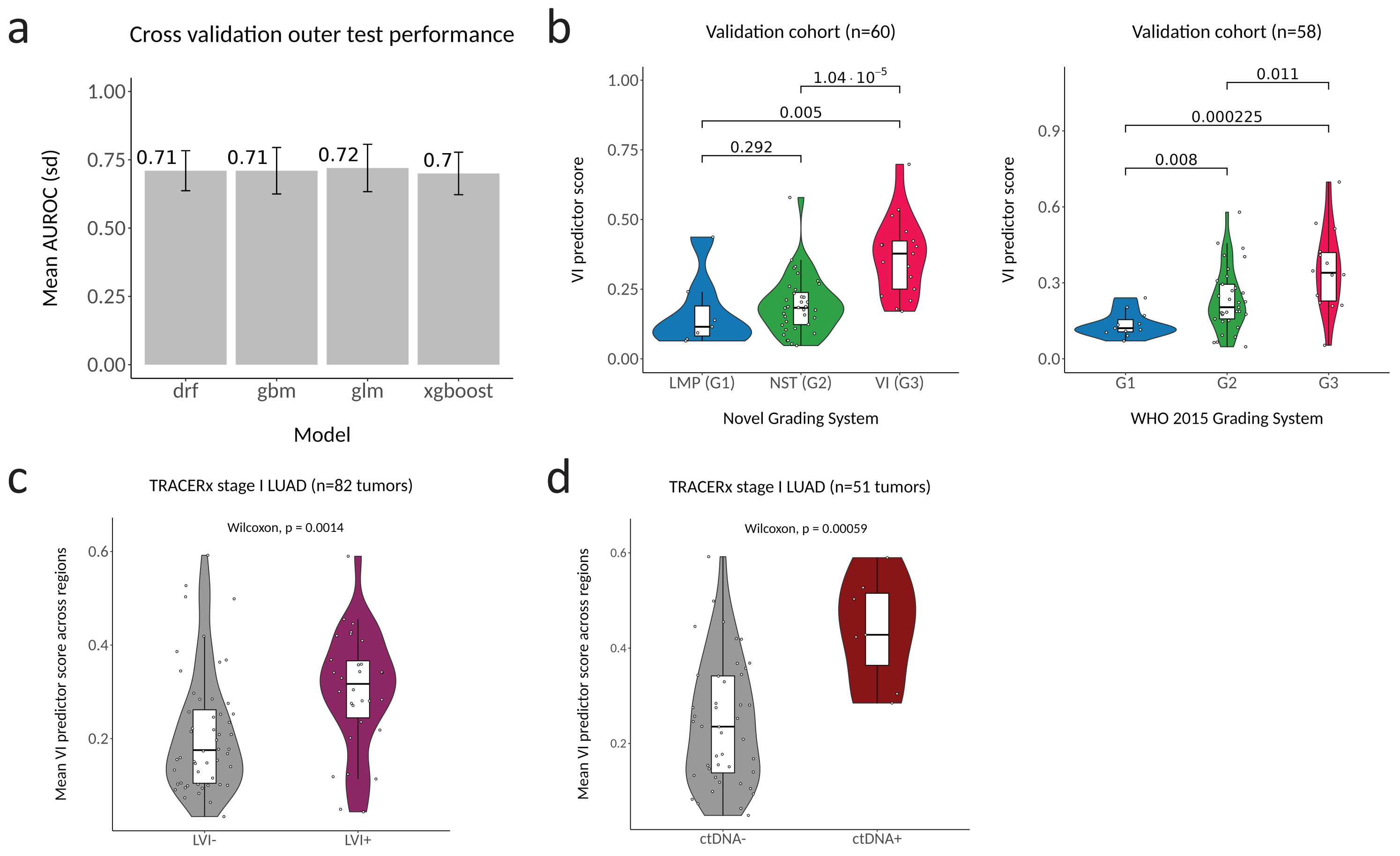


**Extended Data Figure 6. VI predictor development and validation.**
