## Extended Data Fig. 2 for "Identification of a gene expression signature of vascular invasion and recurrence in stage I lung adenocarcinoma via bulk and spatial transcriptomics"

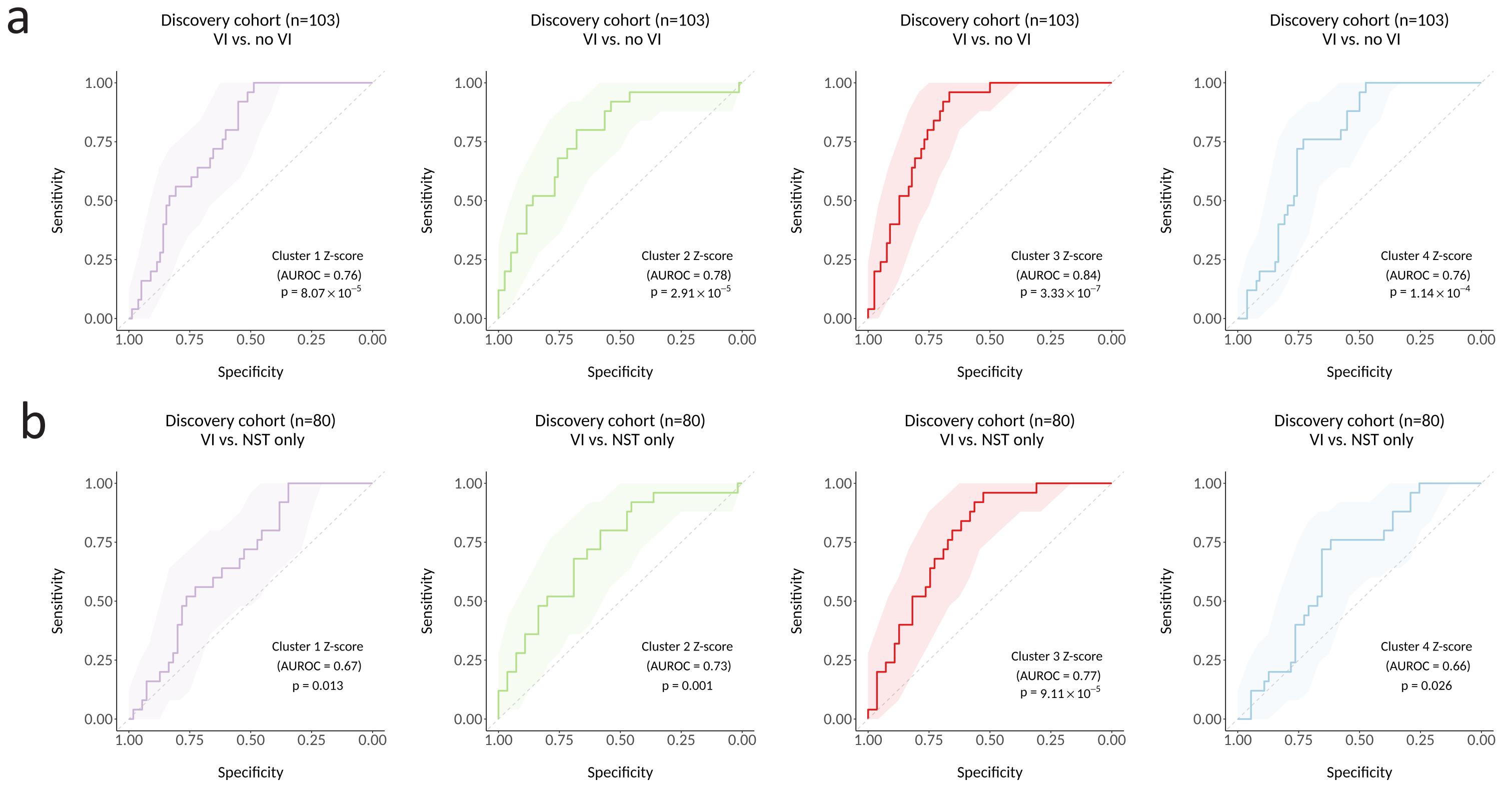


**Extended Data Fig. 2. The four VI gene expression clusters individually predict VI in the discovery cohort even in the absence of LMP tumors.**
